# NicheCorr: Identifying Niche-Specific Differential Gene Co-Expression in Spatial Transcriptomics

**DOI:** 10.64898/2026.10.23.761453

**Authors:** Shuyi Yang, Yen-Yi Ho, Daping Fan

## Abstract

Cell–cell communication is central to biological processes such as immune responses and tumor progression, yet the role of spatial proximity between distinct cell types in shaping gene co-expression remains poorly understood. Existing approaches primarily identify differences in average gene expression across tissue regions, often overlooking how local cellular microenvironments reshape gene–gene interactions within cells. To address this gap, we introduce NicheCorr, a statistical procedure that identifies cell type–specific niche-associated gene co-expression changes within distinct spatial niches. In simulations, NicheCorr attained the highest power for detecting niche-dependent co-expression. It was also the only method evaluated that maintained Type I error control when mean expression varied nonlinearly with the niche, a setting in which CS-CORE, scHOT without mean adjustment, and the Pearson test all exhibited inflated false positive rates. Furthermore, NicheCorr integrates ligand–receptor (LR) inference, enabling a comprehensive view of spatially structured signaling interactions. We applied the framework to the CosMx SMI Human Liver FFPE Dataset and the Xenium Human Pancreatic Cancer Dataset, identifying 52 gene pairs in liver tumor cells whose co-expression changes with T-cell niche composition and 48 gene pairs in pancreatic cancer whose fibroblast niche–dependent co-expression differs between tumor and CFTR^−^ tumor cells, and uncovering potential LR interactions that may underlie these differential co-expression patterns.

## Introduction

Cell–cell communication is fundamental to biological processes, shaping cellular development, immune responses, disease progression, and tumor dynamics within the tissue microenvironment. These interactions are driven both by intrinsic transcriptional programs within individual cells and by extrinsic cues, including paracrine signals from neighboring cells. In particular, the spatial proximity of interacting cells may influence gene co-expression patterns in the cells of interest. For example, tumor cells adjacent to immune cells may exhibit distinct gene coexpression patterns from tumor cells located farther from immune cells. Characterizing how such spatial cues shape gene co-expression is crucial for elucidating mechanisms of intercellular communication. Advances in single-cell RNA sequencing (scRNA-seq) and spatial transcriptomics enable high-throughput, spatially resolved gene-expression profiling, facilitating the study of cell–cell interactions across cell types and the inference of ligand–receptor signaling.

Despite these advances, current methods for studying cell–cell communication have important limitations. Many approaches focus exclusively on differences in mean gene expression. For example, Niche-DE^1^ identifies genes whose average expression varies across spatial niches but does not assess whether gene–gene relationships change with spatial proximity. Yet cell–cell signaling may alter how genes co-vary, rather than merely changing their expression levels.

Methods that model gene co-expression also have important limitations. SpaceBF^2^ estimates spatially varying linear associations but does not directly capture nonlinear changes in co-expression induced by the local cellular niche. spMOCA^3^ estimates a global gene–gene covariance structure across the tissue using a matrix-normal model, thereby overlooking variation across local microenvironments. SpatialCorr^4^ characterizes spatially varying correlation structures and tests whether correlations differ across or within regions, but does not model continuous changes in gene relationships as niche cell-type composition varies. CNNC^5^ uses supervised deep learning to predict gene interactions from expression data, but incorporates neither spatial context nor niche-dependent co-expression dynamics.

Overall, existing approaches either focus on mean expression, assume a single global gene network, test for correlation changes without explicitly modeling niche effects, or ignore spatial context altogether. None directly model how gene co-expression varies with the spatial proximity of interacting cell types within local microenvironmental niches.

To address this gap, we develop NicheCorr, a copula-based computational framework built on generalized additive models for location, scale, and shape (GAMLSS)^6^ to characterize how spatial proximity shapes gene co-expression across interacting cell types. Unlike Niche-DE^1^, which focuses on mean expression, NicheCorr models nonlinear, niche-associated changes in gene–gene correlations while also quantifying corresponding changes in mean expression. Within NicheCorr, kernel-weighted physical distances between interacting cell types, or coordinates within specific microenvironments, are treated as continuous covariates. The model then evaluates whether the copula dependency parameter changes as cells become closer to or farther from one another. By integrating ligand–receptor inference methods, including NicheNet^7^ and Niche-LR, NicheCorr provides a comprehensive characterization of cell–cell communication that extends beyond mean-expression analysis to reveal higher-order, spatially structured interactions.

The framework yields naturally interpretable and generalizable findings. For example, it can determine whether genes *g*_1_ and *g*_2_ are more or less co-expressed in tumor cells (index cells) when T cells (niche cells) are nearby; how this relationship changes with the local abundance of niche cells; whether specific ligand–receptor interactions explain co-expression differences across neighboring cell types; and whether these effects vary among tumor-cell subtypes. NicheCorr can further assess whether co-expression between *g*_1_ and *g*_2_ in tumor cells is associated with expression of a third gene, *g*_3_, in adjacent niche cells.

This capability is particularly relevant because intercellular signaling is often mediated by specific molecules, such as ligands and cytokines, expressed by neighboring cells. The downstream effects of these mediators on gene co-expression cannot be captured through pairwise analysis alone. By incorporating expression of a candidate mediator gene in niche cells, NicheCorr can test for co-expression changes associated with specific molecular signals. To our knowledge, no existing framework models how gene co-expression within one cell type varies with expression of a third gene in a spatially adjacent cell type. These capabilities make NicheCorr well suited to dissecting how microenvironmental context shapes gene regulatory relationships across interacting cell types.

## Results

### Overview of NicheCorr

Gene co-expression is typically estimated as a single sample-level quantity that averages over variation across cells.^8,9^ We developed NicheCorr, a framework that models gene–gene correlation as a function of cell-level covariates — the composition of a cell’s spatial neighborhood, the expression of a third gene, or other cell-level features — enabling co-expression to be characterized as it varies across a tissue.

#### General model

Given a gene expression count matrix **Y** ∈ℝ^*N×G*^ for *N* cells and *G* genes, with entry *Y*_*i,g*_ denoting the count of gene *g* in cell *i*, let **x**_*i*_ = (*x*_*i*1_, … , *x*_*iK*_)^*⊤*^ denote a vector of *K* cell-level covariates for cell *i* (index cell). These covariates may describe biologically meaningful cellular contexts, including spatial microenvironment composition, developmental pseudo-time, average gene expression in neighboring cells, or experimental condition. For a gene pair (*g*_1_, *g*_2_) in the index cell, let *ρ*_*i*_(*g*_1_, *g*_2_) denote their cell-specific correlation. We model its Fisher-transformed value, *z*_*i*_(*g*_1_, *g*_2_), as a smooth function of these covariates:

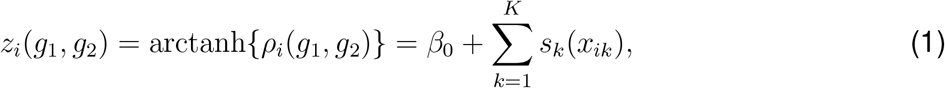

where *β*_0_ is an intercept and *s*_*k*_(·) is a smooth function that captures the nonlinear association between covariate *k* and gene co-expression. The subscript *i* indicates that *z*_*i*_(*g*_1_, *g*_2_) and *ρ*_*i*_(*g*_1_, *g*_2_) are evaluated at the covariate values **x**_*i*_ of cell *i*. The estimated correlation is obtained by applying the inverse Fisher transformation,

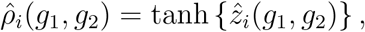

which constrains it to (−1, 1). This additive formulation yields correlation curves for each covariate, enabling direct comparison of how cellular contexts regulate gene-gene co-expression. Gene pairs with statistically significant covariate-dependent correlations are assembled into a co-expression network, where nodes represent genes and edges represent significant context-dependent relationships.

#### Effective niche as a covariate

In this paper, we take **x**_*i*_ to be the effective niche composition surrounding each cell, adopting the niche-construction approach from Niche-DE^1^. The workflow is illustrated in Figure 1. In Figure 1, for each index cell *i* (*i*∈ *{*1, 2, 3, … , *N}*), we define a local spatial niche using a Gaussian kernel with bandwidth *σ*, which controls the spatial scale of the neighborhood. The abundance of cell type *t* (niche cell type *t*) in this neighborhood is counted and weighted based on their spatial distance from the index cell and by the kernel bandwidth *σ*. When *σ* is small, only nearby cells contribute meaningfully, yielding a highly localized niche; when *σ* is large, more distant cells are included, producing a broader spatial niche. Finally, the surrounding microenvironment of the index cell *i* is represented as a *T* -dimensional vector whose *t*th entry quantifies the kernel weighted abundance of cell type *t* (niche cell type *t, t* ∈{1, … , *T}*) in its neighborhood. The bandwidth is selected using a cosine-distance-based procedure detailed in Section Bandwidth Selection for Effective Niche Estimation. This yields an effective niche matrix **E** ∈ℝ^*N×T*^ , with **x**_*i*_ = (*E*_*i*,1_, … , *E*_*i,T*_)^*⊤*^ denoting the effective niche composition for cell *i*. We then have

**Figure 1:**
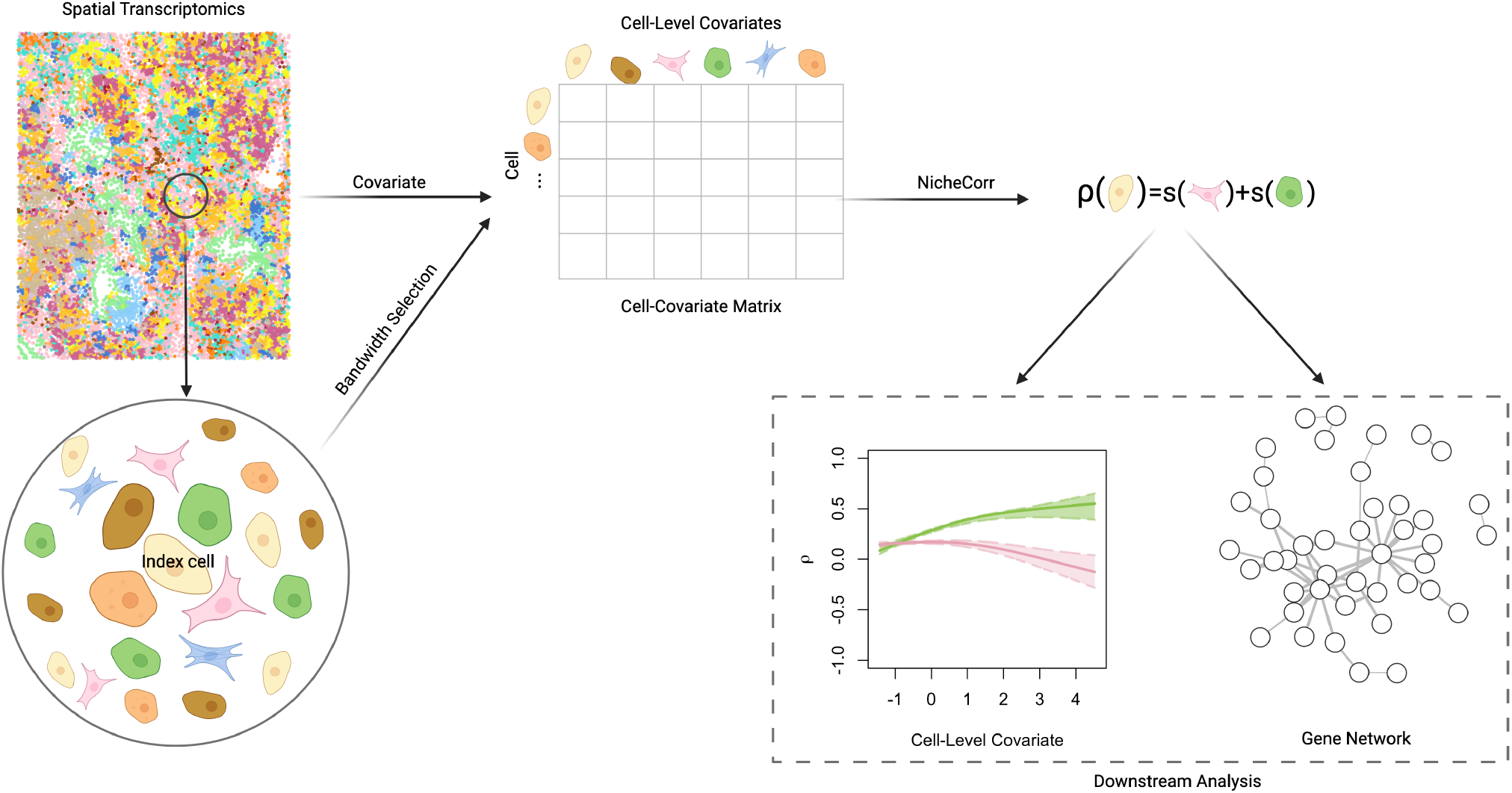
Overview of the NicheCorr framework. (Left) Spatial transcriptomics data with cells annotated by cell type. For each index cell (center), neighboring cells within a kernel bandwidth *σ* contribute to its effective niche, with closer cells receiving greater weights. The bandwidth is selected using a cosine-distance-based procedure. (Middle) The effective niche is summarized as an *N× T* matrix describing the abundance of each cell type in the local neighborhood of every cell. (Right) For each gene pair, co-expression (*ρ*) is modeled as changing with niche composition via additive smooth terms for selected cell types, producing gene–gene correlation curves that enable direct comparison across cell types. Significant niche-dependent gene pairs are assembled into a niche-related gene co-expression network.

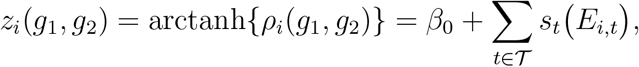

where *T* is the set of cell types of interest, and *E*_*i,t*_ is the effective niche abundance of cell type *t* around cell *i*.

#### Neighboring-cell gene expression as a covariate

The framework also accommodates covariates derived from the gene expression in the niche cells, rather than their cell-type composition. For an index cell *i* and a neighboring cell type *t*, we summarize the local expression of gene *g*_3_ among nearby cells of type *t* using a Gaussian kernel with bandwidth *σ*, denoted 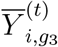 (Equation (6)). Setting *K* = 1 and 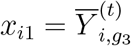 gives

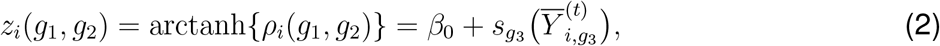

which tests whether the co-expression of genes *g*_1_ and *g*_2_ in the index cell varies with the expression of a third gene, *g*_3_, in surrounding niche cells of type *t*. Conditioning on a third gene enables the framework to link co-expression changes to the activity of specific molecular mediators, such as ligands or cytokines, expressed in neighboring cells.

### NicheCorr provides low-bias, low-RMSE estimation

We conducted a simulation study to evaluate the performance of NicheCorr. We focused on a single niche cell type *t* and, for simplicity, dropped the cell index *i*, denoting effective niche abundance by *E*_*t*_ rather than *E*_*i,t*_. We simulated *E*_*t*_ over a sequence ranging from −0.63 to 4.2 to mimic the range of standardized effective niche values observed in real data (negative values arise because *E*_*t*_ is centered and scaled; see Section Spatial Niche Quantification). We then approximated the distributional parameters *µ*_1_, *µ*_2_, and *ρ* as functions of *E*_*t*_ using polynomial regression up to degree five 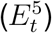, while holding the dispersion parameters *ϕ*_1_ = 1.26 and *ϕ*_2_ = 1.94 fixed. These approximations were based on a real gene pair, APOE and S100A6 identified in our experimental data analysis, to reflect biological relevance.

Using these approximations, we generated paired count data (*y*_1_, *y*_2_) for genes *g*_1_ and *g*_2_, respectively, from a Gaussian copula with negative binomial (NB) marginal distributions:

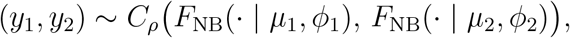

where *C*_*ρ*_ is a Gaussian copula with correlation parameter *ρ, µ*_1_ and *ϕ*_1_ are the mean and dispersion parameters for *y*_1_, *µ*_2_ and *ϕ*_2_ are the mean and dispersion parameters for *y*_2_, and *ρ* measures their dependence. We then fit our proposed model to the simulated data, using *N* = 15,000 observations.

To further investigate the effect of sample size, we performed stratified random sampling at three levels: 80% (*N* = 12,000), 50% (*N* = 7,500), and 30% (*N* = 4,500) of the full simulated dataset. The simulations were repeated *R* = 1,000 times to assess the accuracy of the parameter estimates. Figure 2 shows the results: the red solid lines represent the true parameter functions, the black dashed lines show the mean estimates across the 1,000 iterations, and the shaded areas indicate the pointwise 5%-95% quantile ranges. These ranges cover the true smooth functions in all settings. The left, middle, and right panels correspond to *µ*_1_, *µ*_2_, and *ρ*, respectively.

**Figure 2:**
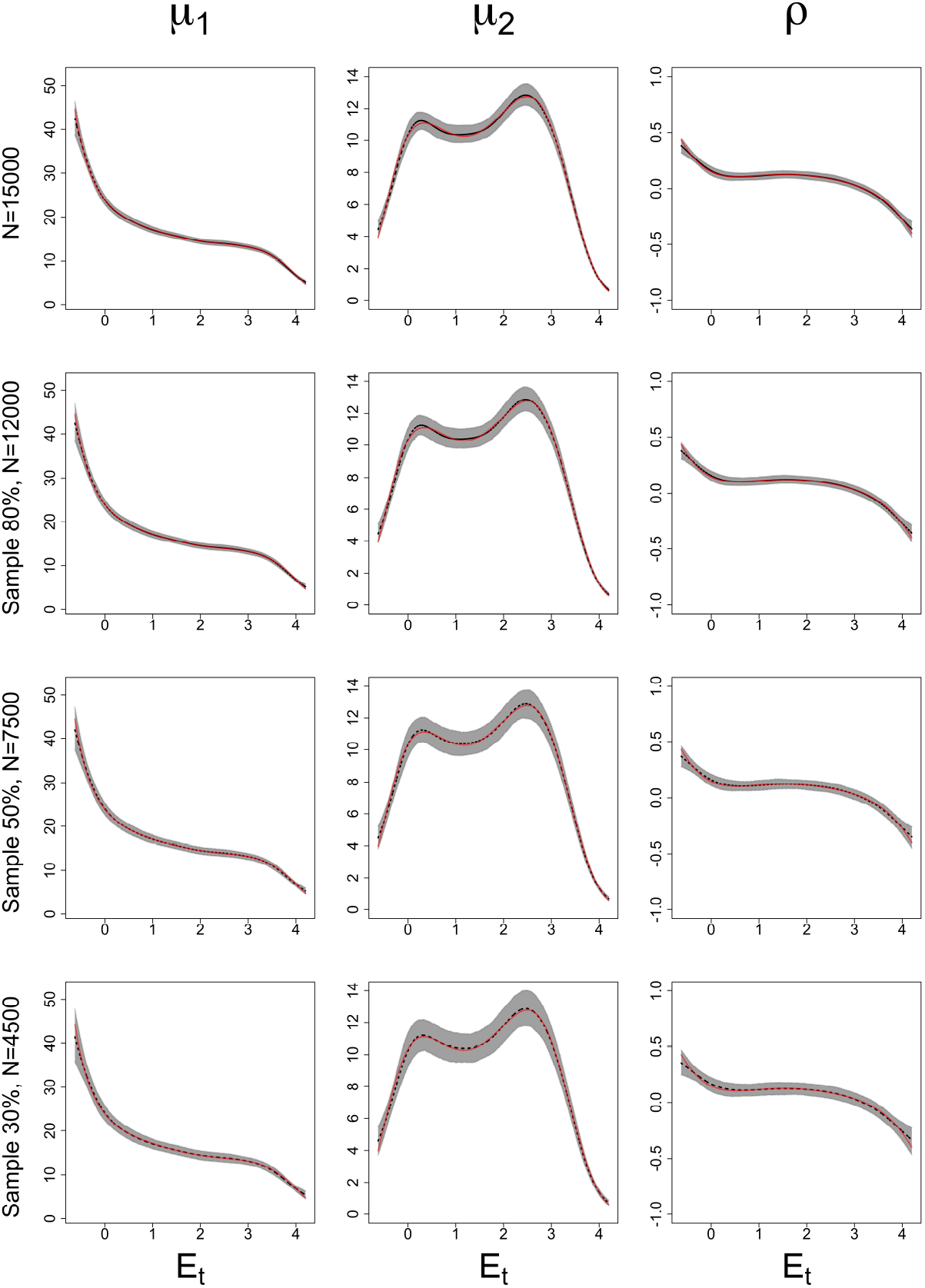
Simulation study results. Red solid lines represent the true smooth functions, and black dashed lines show the mean estimates across 1,000 iterations. Shaded areas indicate the pointwise 5%-95% quantile ranges. The full simulated dataset contains *N* = 15,000 observations, with stratified random subsamples of 80% (*N* = 12,000), 50% (*N* = 7,500), and 30% (*N* = 4,500).

We quantified estimation accuracy by computing the bias and root mean squared error (RMSE) for each parameter at each subsampling level, defined as

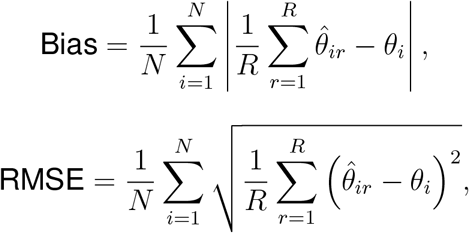

Bias = 1

where *R* = 1,000 is the number of simulation replicates, *θ*_*i*_ is the true parameter value at observation *i*, and 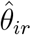 is its estimate in replicate *r*. We additionally computed, for each parameter, the average length of the pointwise 5%-95% quantile interval across the 1,000 simulations. The bias, RMSE, and average 5%-95% quantile interval length for each parameter are reported in Table 1.

**Table 1:** Simulation results for NicheCorr under varying sample sizes. The table shows the average bias, RMSE, and mean length of the pointwise 5%-95% quantile interval for each parameter across 1,000 simulation replicates. Data were generated from a Gaussian copula with NB marginal distributions, where *µ*_1_, *µ*_2_, and *ρ* were modeled as fifth-degree polynomial functions of the effective niche abundance *E*_*t*_. Results are shown for the full dataset (*N* = 15,000) and stratified subsamples of 80%, 50%, and 30%.

|  | Parameter | Bias | RMSE | Avg 5%-95% Quantile Length (%) |
| --- | --- | --- | --- | --- |
| $N = 15,000$ | $\mu_1$ | 0.098 | 0.478 | 1.510 |
| | $\mu_2$ | 0.087 | 0.324 | 0.995 |
| | $\phi_1$ | 0.002 | 0.015 | 0.049 |
| | $\phi_2$ | 0.002 | 0.025 | 0.081 |
| | $\rho$ | 0.005 | 0.023 | 0.072 |
| Sample 80%, $N = 12,000$ | $\mu_1$ | 0.099 | 0.527 | 1.676 |
| | $\mu_2$ | 0.086 | 0.358 | 1.115 |
| | $\phi_1$ | 0.002 | 0.017 | 0.056 |
| | $\phi_2$ | 0.003 | 0.028 | 0.091 |
| | $\rho$ | 0.006 | 0.025 | 0.079 |
| Sample 50%, $N = 7,500$ | $\mu_1$ | 0.120 | 0.643 | 2.028 |
| | $\mu_2$ | 0.080 | 0.432 | 1.363 |
| | $\phi_1$ | 0.002 | 0.021 | 0.068 |
| | $\phi_2$ | 0.005 | 0.035 | 0.116 |
| | $\rho$ | 0.007 | 0.031 | 0.097 |
| Sample 30%, $N = 4,500$ | $\mu_1$ | 0.157 | 0.801 | 2.522 |
| | $\mu_2$ | 0.089 | 0.532 | 1.692 |
| | $\phi_1$ | 0.003 | 0.027 | 0.088 |
| | $\phi_2$ | 0.006 | 0.046 | 0.150 |
| | $\rho$ | 0.008 | 0.039 | 0.122 |

The results show that the full dataset exhibited the narrowest 5%–95% quantile bands and the lowest bias and RMSE values. As the sampling proportion decreased from 80% to 50% to 30%, the quantile bands widened, and both bias and RMSE increased, while overall coverage rates remained similar across sample sizes. The average runtime per simulation replicate was 7.115, 5.605, 3.379, and 1.964 minutes for the full dataset and the 80%, 50%, and 30% sampling levels, respectively.

## NicheCorr controls the false positive rate

To evaluate Type I error control, we conducted simulation studies comparing NicheCorr with several existing approaches, including CS-CORE^9^, scHOT with and without *µ* estimation^8^, the Pearson correlation test, and the Binned Pearson Permutation Test. We simulated gene expression data for 5,000 cells and 50 genes under a null scenario in which gene pairs were uncorrelated. For each cell, we assigned an effective niche covariate, *E*_*t*_, from a fixed sequence ranging from −0.63 to 4.2, representing the spatial niche abundance of a neighboring cell type. For each gene, we modeled the mean of the NB distribution as a smooth function of the effective niche, *s*_*µ*_(*E*_*t*_), using the following sine function :

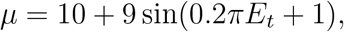

and set the dispersion parameter to *ϕ* = 0.9. Gene expression counts were simulated from a Gaussian copula model with NB marginal distributions and an identity correlation matrix.

We applied NicheCorr and the comparison methods to test gene-gene correlations across all 1,225 gene pairs and collected the resulting *p*-values under the null hypothesis of no correlation between genes. For the Binned Pearson Permutation Test, we divided *E*_*t*_ into 30 bins and computed Pearson correlations within each bin. We then applied the Fisher *z*-transformation to the bin-specific correlations and used their dispersion across bins as the test statistic. Statistical significance was assessed by permuting *E*_*t*_ across cells, recomputing the bin-specific correlations, and comparing the observed dispersion with its permutation distribution based on 1,000 permutations. Details of the procedure and bin selection are provided in the Supplementary Materials.

scHOT tests whether pairwise gene-gene correlations change along a continuous trajectory, here represented by the effective niche covariate *E*_*t*_. A key limitation is that scHOT does not explicitly model changes in gene expression means. When applied without *µ* estimation, the method operates directly on raw counts; therefore, detected signals may reflect both dynamic co-expression and joint shifts in marginal means across *E*_*t*_. In contrast, with *µ* estimation, mean expression is first modeled along *E*_*t*_ using a generalized additive model (GAM), and correlation is then assessed using the adjusted data. This approach aims to detect correlation changes beyond marginal mean trends. We implemented CS-CORE^9^ with the default values.

To evaluate false-positive control, we compared the resulting *p*-value distributions with the expected Uniform(0, 1) distribution using quantile-quantile (Q-Q) plots, shown in Figure 3a. Among all methods, NicheCorr exhibited the closest alignment with the *y* = *x* line, indicating the best Type I error control. scHOT with *µ* estimation also aligned closely with the theoretical distribution, whereas scHOT without *µ* estimation showed moderate deviation below the diagonal, suggesting mild false-positive inflation. CS-CORE produced *p*-values smaller than expected under the uniform distribution, yielding an inflated false-positive rate. In addition, CS-CORE assumes strictly positive cell sequencing depths, which limits its applicability to datasets with small gene panels. The Binned Pearson Permutation Test showed slightly greater variability in the upper tail, whereas the standard Pearson correlation test performed the worst, with a strong accumulation of extremely small *p*-values near zero, reflecting severe Type I error inflation under the null scenario.

**Figure 3:**
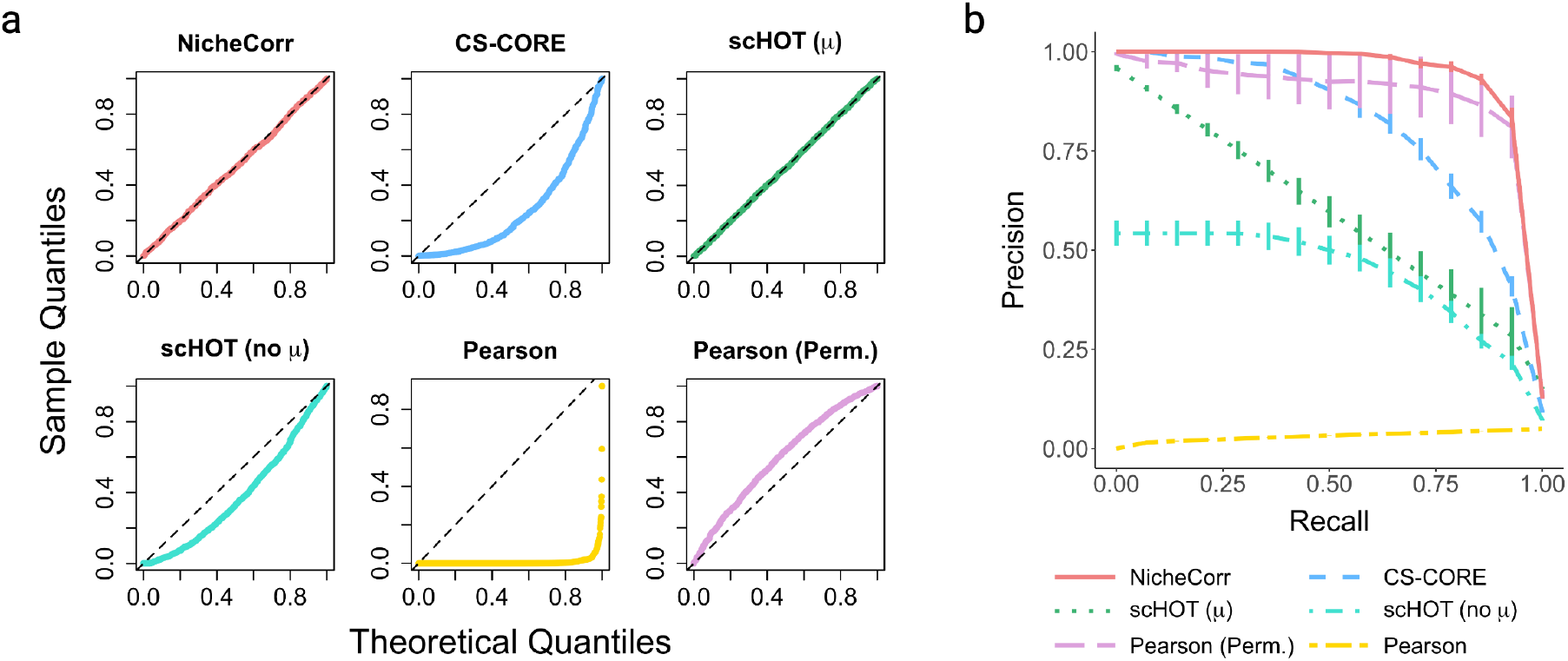
**a** Q-Q plots for NicheCorr, CS-CORE, scHOT with and without *µ* estimation, the Pearson correlation test, and the Binned Pearson Permutation Test (Pearson (Perm.)). The plot compares empirical *p*-value distributions with the expected Uniform(0, 1) distribution under the null hypothesis of no gene-gene co-expression. The black dashed line represents the theoretical *y* = *x* line; closer alignment indicates better Type I error control. NicheCorr and scHOT with *µ* estimation closely follow the diagonal, indicating good false-positive control. Pearson (Perm.) shows mild deviation, while scHOT without *µ* estimation departs more noticeably from uniformity. CS-CORE exhibits substantial deviation, with an excess of small *p*-values, and the standard Pearson test shows a strong bias toward small *p*-values. **b** Precision-recall curves for NicheCorr, CS-CORE, scHOT with and without *µ* estimation, the Pearson correlation test, and the Binned Pearson Permutation Test (Pearson (Perm.)) under the dynamic-correlation simulation design. 5% of gene pairs exhibit co-expression, with all remaining pairs having *ρ* = 0. Curves show the mean across 10 simulations; vertical bars indicate the standard error across the 10 simulation replicates. NicheCorr achieves the highest precision across the evaluated recall levels, yielding the largest area under the precision-recall curve.

### NicheCorr achieves higher statistical power/precision-recall

Having evaluated false-positive control under the null scenario, we next assessed the power of each method under an alternative scenario in which correlations between gene pairs vary with the effective niche abundance *E*_*i,t*_. We simulated expression data for 50 genes across 5,000 cells from a Gaussian copula distribution with NB marginal distributions. We defined an index set *D* comprising 5% of all gene pairs as truly co-expressed. For each cell *i*, the pairwise correlation between genes *g*_1_ and *g*_2_ was defined as

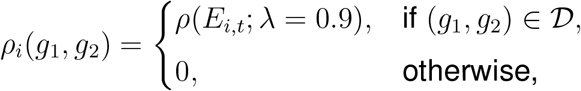

where *λ* is a scaling factor that controls the strength of the correlation signal. The correlation function was defined as

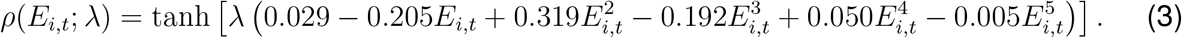

Thus, the set of gene pairs with nonzero correlations was identical across cells, while the correlation values varied smoothly with *E*_*i,t*_. Counts were then generated using this correlation structure, with a constant overdispersion parameter.

We repeated the entire simulation 10 times. For each method, we plotted the precision-recall (PR) curve, as shown in Figure 3b. Each PR curve represents the mean across the 10 replicates, evaluated on a grid of 10 recall levels; vertical bars indicate the standard errors across replicates. NicheCorr performs the best: precision remained near 1 across most of the recall range and decreased only when recall approached 1, indicating accurate detection of gene pairs with dynamic co-expression. The Binned Pearson Permutation Test exhibited comparable performance, with precision remaining high across most recall levels but showing greater variability at high recall. CS-CORE was precise at low recall, but its precision gradually decreased as recall increased.

Both scHOT variants performed worse than NicheCorr and CS-CORE, indicating lower sensitivity to dynamic correlation patterns. scHOT with *µ* estimation showed substantially lower precision, with a steady decline as recall increased, while scHOT without *µ* estimation performed even worse, with low precision even at low recall levels. The standard Pearson correlation test performed poorly overall, with precision remaining close to zero across the recall range, indicating an inability to distinguish true signals from noise under this niche-varying correlation scenario. Overall, these results demonstrate that NicheCorr provides a favorable balance between precision and recall when gene–gene correlations vary smoothly with the effective niche, outperforming competing approaches.

We evaluated the statistical power of NicheCorr through simulation studies and compared its performance with that of existing methods: scHOT^8^ with and without *µ* estimation, CS-CORE^9^, the Pearson correlation test, and the Binned Pearson Permutation Test. The simulations were designed to mimic realistic data in which gene-gene correlations vary dynamically with a continuous, cell-type-specific effective niche.

We simulated expression counts for 5,000 cells and 50 genes from a Gaussian copula distribution with NB marginal distributions. The correlation structure between genes was specified by a smooth function of the effective niche, *E*_*t*_. This function included a scaling factor, *λ*, evaluated at 27 equally spaced values ranging from 0 to 1.3. When *λ* = 0, the correlation curve equals zero, representing a null case with no dynamic co-expression. As *λ* increases, the correlation curves deviate increasingly from zero, producing stronger niche-dependent correlations across *E*_*t*_. Correlation curves for different values of *λ* are shown in Figure 4a, and the correlation function is given in Equation (3).

**Figure 4:**
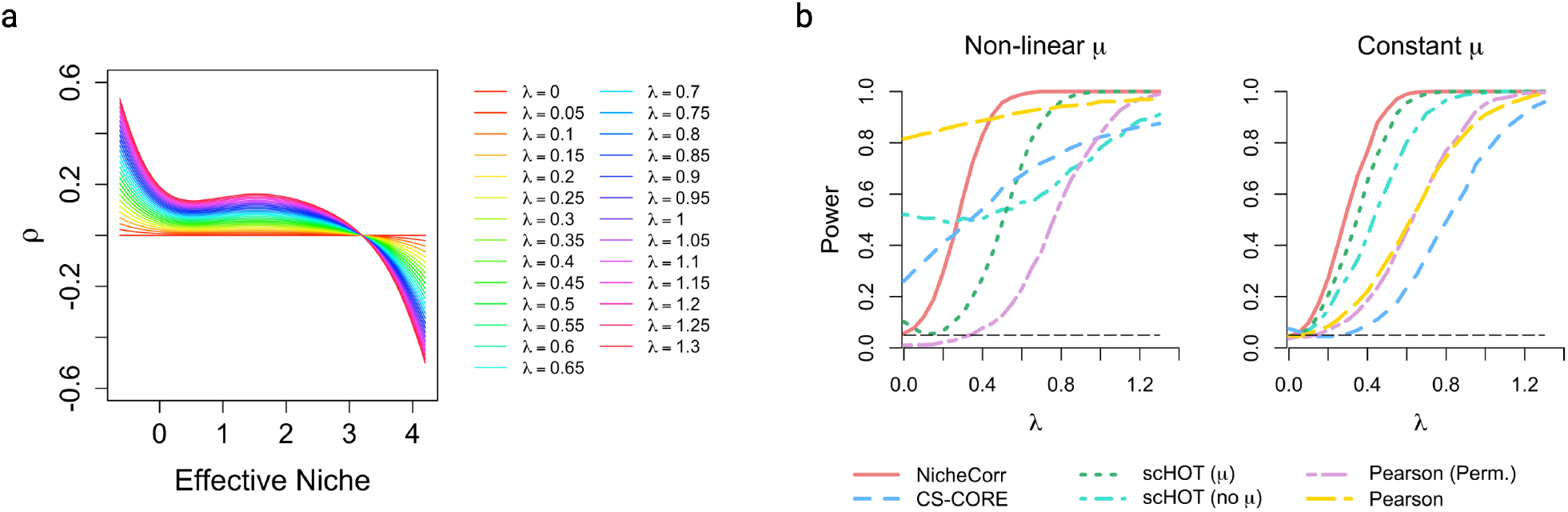
**a** Correlation function *ρ*(*E*_*t*_) used in the simulations for different values of the scaling factor *λ*, which controls the magnitude of departure from zero. Each colored curve shows gene-gene correlation along *E*_*t*_ for a different value of *λ*. When *λ* = 0, there is no dynamic correlation; larger values of *λ* produce stronger correlation dynamics. **b** Statistical power to detect *E*_*t*_-dependent correlation as a function of *λ* across methods. Two mean structures were considered: (i) *µ* simulated from a uniform distribution (constant-mean setting) and (ii) *µ* derived from a real gene pair (nonlinear-mean setting). Power was defined as Power 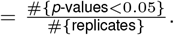. The horizontal black dashed line marks the significance level of *α* = 0.05. NicheCorr consistently achieved the highest power across all values of *λ* , followed by scHOT with *µ* estimation. Pearson-based methods and CS-CORE showed limited ability to detect dynamic correlation.

For each simulated cell, gene-gene correlations were modeled using a copula distribution with a correlation matrix derived from *ρ*(*E*_*t*_; *λ*). Gene counts were generated from NB distributions with constant dispersion and under two distinct mean structures to assess robustness to different expression patterns. In the first setting (nonlinear *µ*), the mean parameters were specified as smooth, nonlinear functions of the effective niche *E*_*t*_, mimicking realistic gene expression dynamics observed for the HSPB1-SELENOP gene pair. This setting reflects biologically plausible scenarios in which gene expression levels vary continuously with cellular context.

In the second setting (constant *µ*), mean expression levels were constant and independent of *E*_*t*_. This design isolates the effect of dynamic correlation from marginal mean changes, ensuring that any detected signal arises solely from *E*_*t*_-dependent correlation rather than shifts in gene expression levels. This simulation design captures the overdispersed count distribution and zeroinflated counts commonly observed in scRNA-seq data.

CS-CORE was applied globally to estimate a full gene–gene correlation matrix and subsequently test the correlation between each gene pair of interest. All other methods were applied directly to the raw count data for each gene pair of interest to model dynamic co-expression patterns along *E*_*t*_.

For each value of *λ*, 1,000 datasets were simulated. Each method produced a *p*-value for testing the null hypothesis of no dynamic correlation. Statistical power was estimated as the proportion of replicates in which the null hypothesis was rejected at the 5% significance level. The power results are shown in Figure 4b.

Across both the nonlinear and constant *µ* settings, NicheCorr consistently achieved the highest statistical power. In the constant-*µ* setting, under the null scenario (*λ* = 0), all methods except for CS-CORE maintained proper Type I error control at approximately 0.05. As the correlation strength *λ* increased, NicheCorr exhibited the fastest increase in power, detecting dynamic co-expression at lower signal strengths than competing approaches. It reached 80% power at moderate signal levels (*λ* = 0.4) and approximately 95% power by *λ* = 0.55.

scHOT ranked second in performance, with power increasing more gradually than NicheCorr and requiring stronger correlation signals to achieve comparable sensitivity. Its performance improved when *µ* was estimated. The Binned Pearson Permutation Test required substantially stronger correlations to reach similar power, while the standard Pearson test performed worse overall. CS-CORE exhibited the weakest performance, requiring a strong signal (*λ* = 1.3) to reach 95% power.

In the nonlinear-*µ* setting, only NicheCorr maintained appropriate Type I error control under the null. In contrast, CS-CORE, scHOT without *µ* estimation, and the Pearson test displayed inflated rejection rates at *λ* = 0, indicating false-positive detections driven by nonlinear mean trends rather than true dynamic correlation.

Although scHOT with *µ* estimation mitigated this inflation, its power increased more slowly, requiring stronger correlation signals (*λ >* 0.65) to exceed 80% power. The Binned Pearson Permutation Test similarly required substantially stronger correlations to achieve comparable sensitivity.

Overall, these results demonstrate that methods that do not explicitly account for dynamic mean structures can produce false-positive detections in the presence of nonlinear mean expression trends. In contrast, NicheCorr robustly controls Type I error while achieving substantially higher power to detect nonlinear, trajectory-dependent gene-gene co-expression.

### CosMx SMI Human Liver Data Analysis

#### NicheCorr identified significant co-expressed gene pairs from CosMx SMI human hepatocellular carcinoma data

We applied NicheCorr to the CosMx SMI Human Liver FFPE spatial transcriptomics dataset^10^ to investigate how the spatial proximity of T cells influences gene co-expression patterns in tumor cells. We treated tumor cells as the index cell type and T cells as the niche cell type. For notational simplicity, we drop the cell index *i* and denote the effective niche abundance of T cells by *E*_*t*_. For each tumor cell, we computed a tumor-T-cell effective niche using Niche-DE^1^, yielding a continuous covariate *E*_*t*_ that quantifies local T-cell proximity.

To ensure sufficient information for correlation estimation in scRNA-seq data with zero inflation, we retained genes with fewer than 50% zero counts among tumor cells. This resulted in 119 genes, yielding 7,021 tumor-cell gene pairs for analysis.

We evaluated each gene pair for evidence that its correlation varied with the effective niche. Spatial transcriptomics data often exhibit global spatial autocorrelation due to broad tissue architecture and biological processes, which can confound the detection of local niche-dependent co-expression patterns. To adjust for this global autocorrelation structure, we first included three Moran’s eigenvector maps (MEMs) as spatial covariates in the marginal predictors. The MEMs capture broad spatial trends, reducing their influence on the estimated niche-dependent effects. We then applied NicheCorr to identify local changes in gene co-expression associated with effective niche composition. This approach allows NicheCorr to focus on local correlation changes attributable to niche composition rather than global spatial trends. The number of MEMs was selected based on the eigenvalues and an elbow plot.

For each gene pair, we compared a null model that assumes a constant correlation, *ρ*(*E*_*t*_) = *ρ*_0_, with a full model that allows the correlation to vary nonlinearly with the effective niche, arctanh {*ρ*(*E*_*t*_)} = *β*_0_ + *s*_*t*_(*E*_*t*_). The models were compared using the Akaike information criterion (AIC). We kept gene pairs with a positive AIC difference, defined as ΔAIC = AIC_null_ −AIC_full_, indicating improved fit of the full model. Statistical significance was assessed using NicheCorr’s hypothesis test for nonconstant correlation along *E*_*t*_. Multiple testing across gene pairs was controlled using the Benjamini-Hochberg (BH) false discovery rate (FDR) procedure, with an adjusted *p*-value threshold of 0.05.

We visualized the results as a volcano-style plot of −log_10_(*p*_adjusted_) against log_10_(ΔAIC) (Figure 5a). Gene pairs exceeding log_10_(ΔAIC) = 2.5, the elbow of the ΔAIC distribution, were prioritized as those that were both statistically significant and showed a substantial improvement in model fit. This procedure identified 52 gene pairs whose correlations varied significantly along the tumor-T-cell niche, as shown in Table 2. Figure 5b illustrates one such pair, HSPB1 and SELENOP. NicheCorr estimated a low or near-zero correlation between these genes when T cells were close to tumor cells, with the correlation increasing as T cells became more distant. These results suggest that spatial proximity to T cells may influence gene-gene relationships within the tumor microenvironment.

**Table 2:**
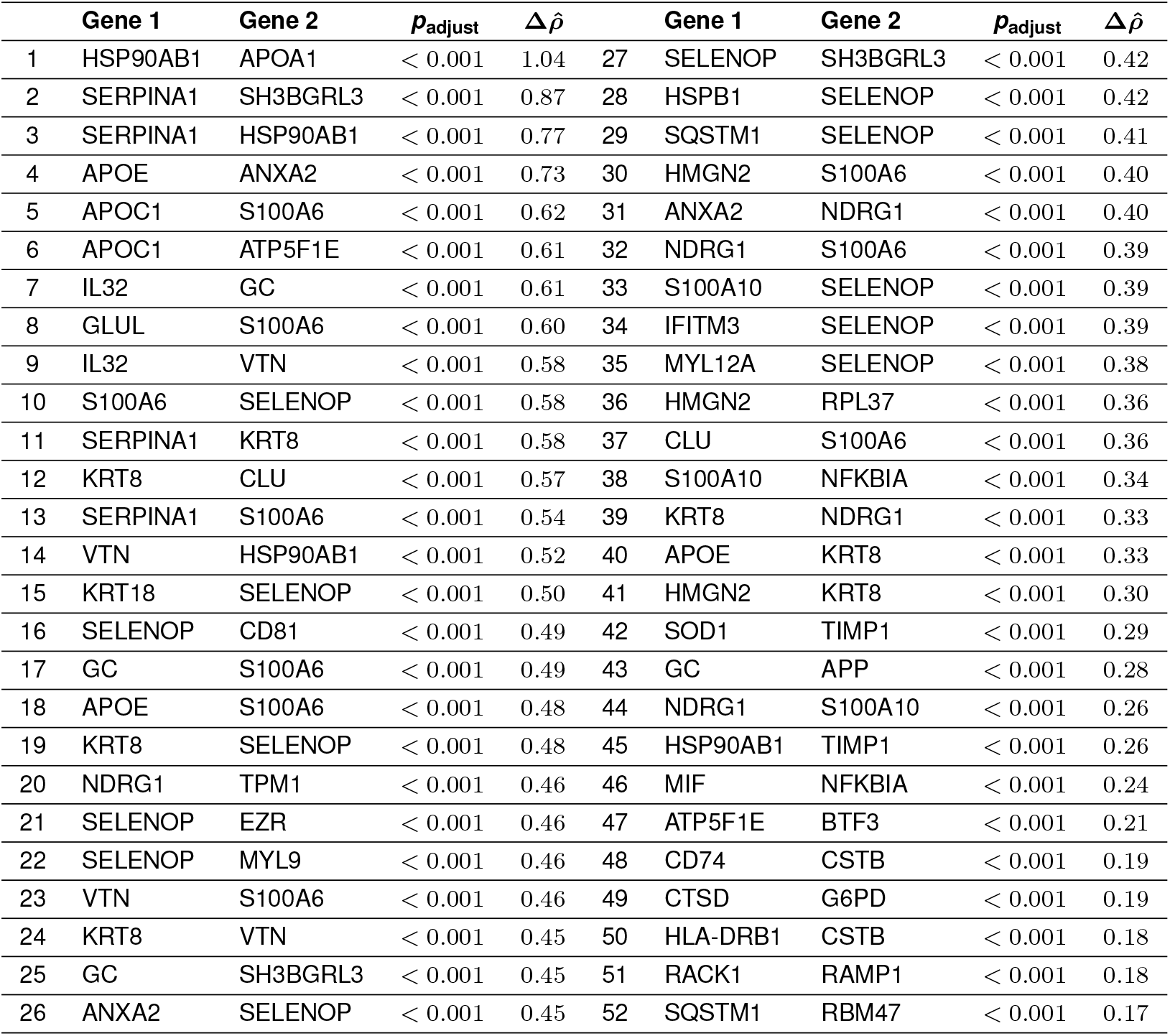
The 52 significant gene pairs in tumor cells with *p*_adjusted_ *<* 0.05 and log_10_(ΔAIC) *>* 2.5, ranked by 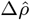. Here, 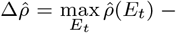 min 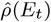denotes the overall range of the estimated correlation across *E*_*t*_. Larger 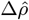 values indicate stronger niche-dependent co-expression. *p*_adjusted_ is the BH-adjusted *p*-value for testing nonconstant correlation along the tumor-T-cell effective niche *E*_*t*_.

**Figure 5:**
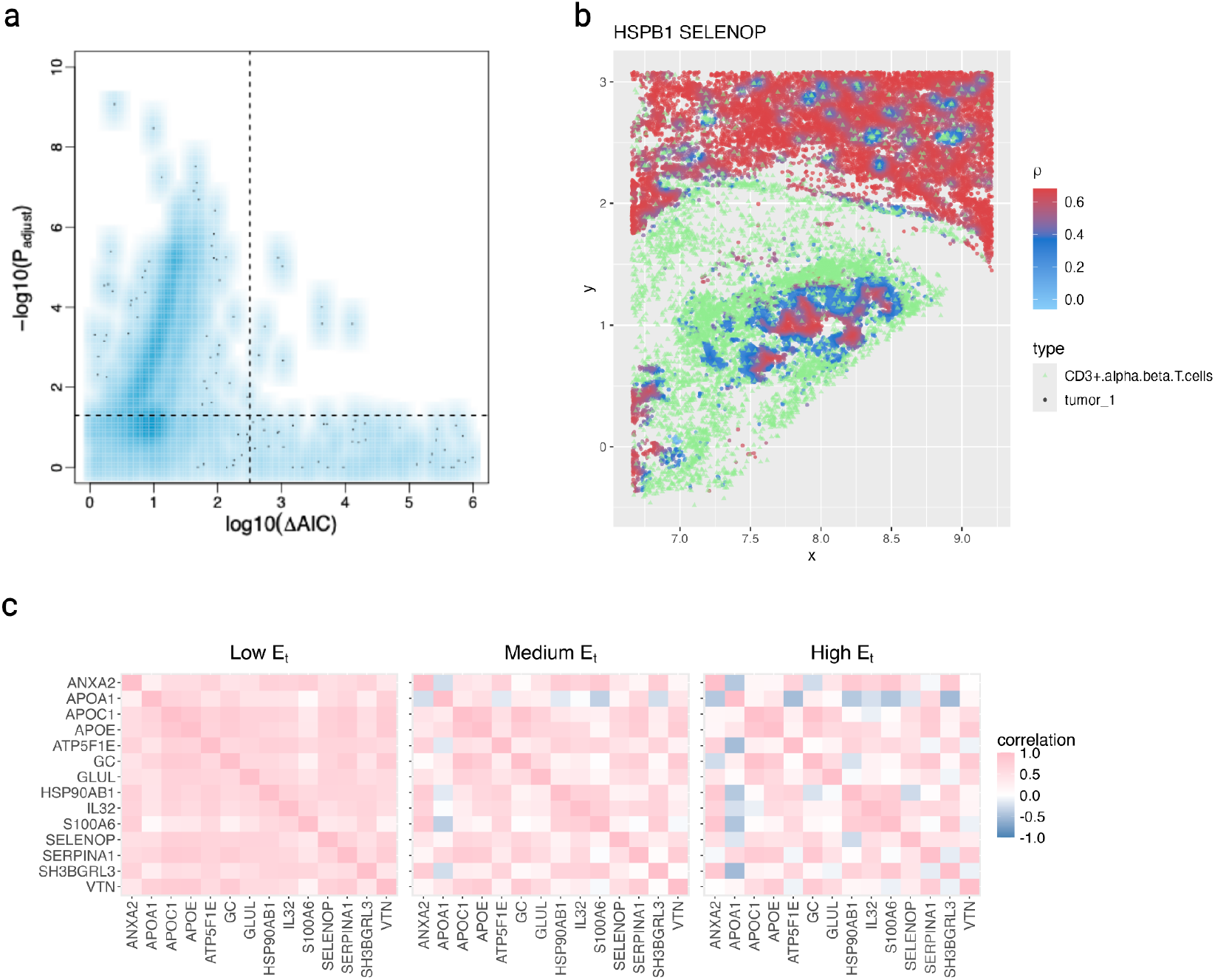
**a** Volcano plot of gene pair correlations. Each point represents a gene pair. The x-axis shows log_10_(ΔAIC), where larger values indicate improved fit of the nonlinear correlation model relative to the null model. The y-axis shows −log_10_(*p*_adjusted_), where *p*_adjusted_ is the BH-adjusted *p*-value for testing whether *ρ*(*E*_*t*_) is nonconstant. The dashed horizontal line marks the adjusted *p*-value threshold of 0.05, and the dashed vertical line marks the ΔAIC threshold. Gene pairs in the upper right region were identified as significant pairs with correlations that vary along the effective niche. **b** Spatial plot of HSPB1-SELENOP correlation. Each point represents a cell at its physical location; circles denote tumor cells, and green triangles denote T cells. The color of each tumor cell reflects the estimated co-expression between HSPB1 and SELENOP at that cell’s location, based on the proximity of T cells in its local niche. The color gradient ranges from blue (*ρ* = −1) to red (*ρ* = 1). **c** *E*_*t*_-stratified correlation heatmaps for the top 10 gene pairs (14 unique genes). Columns show correlations computed within low, medium, and high *E*_*t*_ bands. Color denotes the mean correlation within each band. Gene order is fixed across panels.

We selected the 10 gene pairs involving 14 unique genes with the largest correlation differences across *E*_*t*_. For visualization, tumor cells were partitioned into low, medium, and high *E*_*t*_ bands. Within each band, we computed the average NicheCorr-estimated gene-gene correlation matrix across tumor cells. The resulting heatmaps, shown in Figure 5c, reveal clear nichedependent correlation patterns. The selected pairs include examples in which co-expression decreases with T-cell proximity. In the low *E*_*t*_ band, where T cells are farther from tumor cells, most correlations are strong. As *E*_*t*_ increases, these correlations generally decrease.

### Ligand-receptor analysis

To identify LR interactions associated with dynamic gene co-expression changes captured by NicheCorr, we extended the Niche-LR framework by integrating NicheCorr chi-square test statistics instead of the one-sided *t*-statistics used in Niche-DE.

For each gene pair (*g, g*^*′*^) among the 52 significant pairs involving 43 unique genes, we obtained a chi-square test statistic *χ*^2^(*g, g*^*′*^) from NicheCorr. This statistic tests the entire fitted correlation curve: it quantifies evidence that the cell-specific correlation *ρ*_*i*_(*g, g*^*′*^) varies with the effective niche *E*_*i,t*_ across cells, rather than remaining constant (*H*_0_ : *s*_*ρ*_ = 0). To derive a gene-level statistic, we summed the chi-square statistics across all significant pairs involving a given gene. Thus, genes participating in multiple significant pairs receive larger aggregate test statistics, reflecting stronger evidence of involvement in dynamic co-expression changes.

We visualized the 52 significant gene pairs involving 43 unique genes as an undirected network (Figure 6). Each node represents a gene, and each edge denotes a gene pair whose correlation changes significantly along the tumor-T-cell effective niche, *E*_*t*_. Edge weights are proportional to the log_10_-transformed chi-square test statistic from NicheCorr, reflecting the strength of evidence against constant correlation. To highlight LR associations, nodes are displayed as pie charts. Each colored sector within a node corresponds to a known LR interaction involving that gene. Candidate LR interactions identified using the extended Niche-LR framework are summarized in Table 3, together with the top-ranked gene sets associated with each interaction. The network reveals several hub genes with extensive dynamic co-expression connectivity, including SELENOP, NFKBIA, HSPB1, CLU, ANXA2, NDRG1, APOE, IL32and SOD1. These hubs are centrally positioned and connected to multiple genes, suggesting coordinated regulation within tumor cells that is associated with T-cell proximity.

**Table 3:**
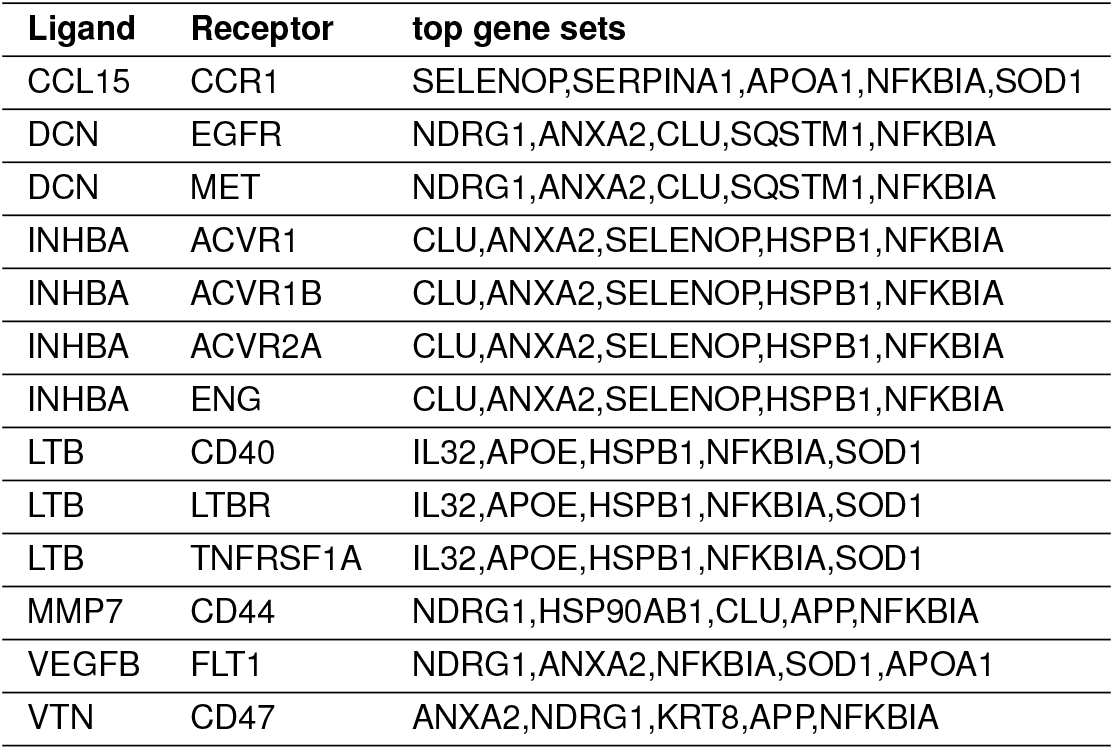
Candidate LR interactions associated with *E*_*t*_-dependent gene co-expression in tumor cells. LR interactions were inferred using the Niche-LR framework extended with NicheCorr gene pair level chi-square statistics. For each LR pair, the top gene set lists the highest-ranked genes associated with the interaction.

**Figure 6:**
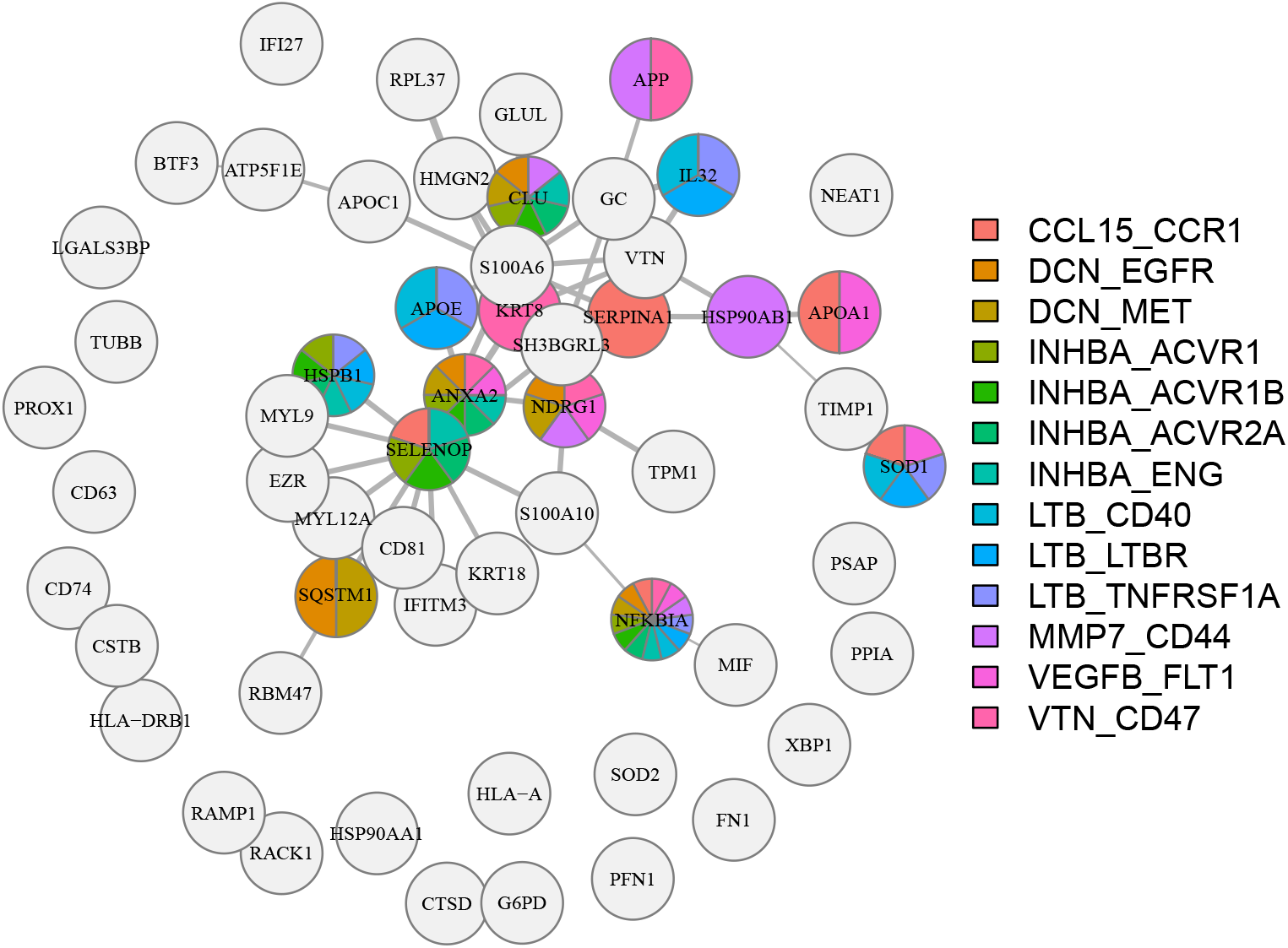
Network of significant gene pairs with *E*_*t*_-dependent co-expression. The 43 gene nodes are connected by edges representing the 52 significant pairs whose correlation, *ρ*(*E*_*t*_), is nonconstant. Edge width is proportional to the log_10_-transformed chi-square test statistic for testing the constancy of *ρ*(*E*_*t*_). Thicker edges indicate stronger evidence of *E*_*t*_-dependent correlation. Genes involved in multiple LR interactions display multiple colored sectors, indicating potentially multifunctional signaling roles.

These results suggest that the local abundance of T cells influences how genes are coexpressed within tumor cells. In particular, genes involved in cell-cell communication, including LR signaling, show clear changes in the strength of their co-expression as T cells become closer to or farther from tumor cells.

Together, these findings indicate that NicheCorr can detect shifts in tumor-cell communication and intracellular gene programs in response to the local immune microenvironment. These shifts reflect not only changes in the expression levels of individual genes, but also changes in how genes interact with one another.

### Pancreatic Cancer Xenium Data Analysis

#### Data preprocessing and quality control

For this study, we analyzed the publicly available Pancreatic Cancer Xenium Human Multi-Tissue and Cancer Panel in situ gene expression dataset from 10x Genomics^11^.

Quality control was performed on the raw count matrix to evaluate overall data quality. For each cell, we calculated the total number of detected transcripts (total counts per cell) and the number of detected genes (genes with nonzero counts). We inspected the distributions of these metrics using violin plots to assess overall data quality and identify potential outliers.

Based on visual inspection of the violin plots, the distributions of total counts and detected genes showed no evidence of low-quality cells or extreme outliers. Therefore, no cell-level filtering was applied, and all cells were retained for downstream analyses. Genes were filtered according to their detection frequency across cells, retaining genes with nonzero counts in at least 10% of cells. This resulted in 149 genes, yielding 11,026 gene pairs for subsequent analyses.

#### Spatial domain identification and selection

To identify biologically meaningful spatial regions within the pancreatic cancer Xenium dataset, we performed spatial domain identification. Spatial domains were defined as groups of cells with similar gene expression profiles that were also spatially proximate within the tissue. This approach integrates transcriptomic information with spatial coordinates, enabling the detection of tissue microenvironment structures that may not be captured by conventional clustering methods based only on gene expression.

Before spatial domain identification, we applied principal component analysis (PCA) to the gene expression matrix to reduce dimensionality. The number of principal components was determined according to the variance explained by each component. We used the first 20 principal components, which captured the major biological variation in the data.

A gene expression-based nearest neighbor graph was constructed using a *k*-nearest-neighbors (KNN) approach, with pairwise distances calculated using Euclidean distance. The number of nearest neighbors was selected using the silhouette score to optimize clustering quality. To model physical proximity between cells, we constructed a spatial neighborhood graph based on cellular spatial coordinates. For each cell, spatial neighbors were defined as cells located within a fixed physical radius. The radius was set to three times the median nearest-neighbor distance, balancing the capture of local spatial structure with sufficient neighborhood connectivity. To jointly incorporate transcriptomic similarity and spatial proximity, the gene expression-based and spatial neighborhood graphs were combined into a single weighted graph, and Leiden clustering was applied to identify spatial domains.

This analysis identified multiple spatial domains across the pancreatic cancer tissue (Figure 7a). To characterize the cellular composition of each domain, we quantified cell-type proportions within each domain (Figure 7c). Based on these compositions, we selected domains 3, 5, 8, which are enriched in tumor cells, CFTR^*−*^ tumor cells, and fibroblasts, also displayed clear spatial organization within the tissue (Figure 7b).

**Figure 7:**
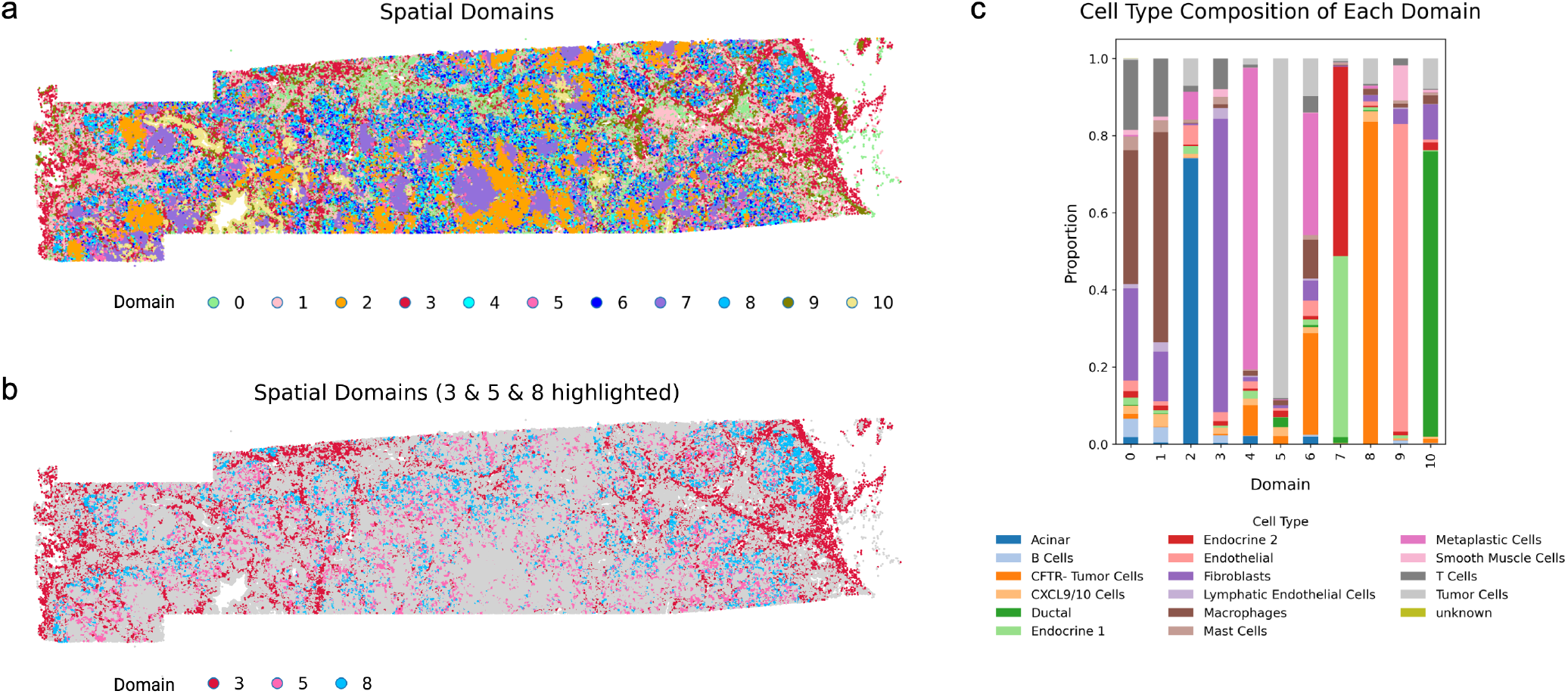
Spatial domain identification and cellular composition in pancreatic cancer tissue. **a** Spatial domains identified by integrating gene expression similarity and spatial proximity using Leiden clustering. Each color represents a distinct domain. **b** Domains enriched in tumor cells (Domain 5), CFTR^*−*^ tumor cells (Domain 8), and fibroblast-associated regions (Domain 3) are highlighted. **c** Cell-type composition of each domain, shown as the proportion of annotated cell types, confirming enrichment of tumor, CFTR^*−*^ tumor, and fibroblast populations in the selected domains.

Using these selected domains, we investigated how gene co-expression patterns in tumor cells and CFTR^*−*^ tumor cells varied with the tumor-fibroblast effective niche and whether these niche-dependent patterns differed between the two tumor groups.

#### Significant Gene Pairs

To investigate whether niche-dependent co-expression patterns differ between tumor cells and CFTR^*−*^ tumor cells, we applied NicheCorr within the selected spatial domains enriched for tumor-fibroblast interactions, including one MEM as a spatial covariate. The effective niche strength, *E*_*t*_, was quantified by the local abundance of fibroblasts surrounding each tumor cell. For each gene pair, we modeled paired gene expression using a bivariate copula framework with NB marginal distributions, allowing the correlation parameter to vary smoothly with *E*_*t*_ and differ between the two tumor groups.

Specifically, the correlation trajectory was modeled as a smooth function of *E*_*t*_ with an interaction term that permits group-specific deviations. This formulation enabled us to test whether correlation curves along niche strength differed between tumor cells and CFTR^*−*^ tumor cells. The null hypothesis states that the two groups share the same correlation trajectory across *E*_*t*_, whereas the alternative allows group-specific correlation dynamics.

Statistical significance was assessed using a chi-square test derived from the smooth interaction term. *p*-values were adjusted across all tested gene pairs using the BH procedure to control the FDR. At an FDR threshold of 0.05, we identified 48 gene pairs with significantly different niche-dependent correlation trajectories between tumor cells and CFTR^*−*^ tumor cells (Table 4).

**Table 4:** The 48 gene pairs with significantly different niche-dependent correlation trajectories between tumor cells and CFTR^*−*^ tumor cells (FDR *<* 0.05), ranked by *p*_adjusted_. *p*_adjusted_ denotes the BH-adjusted *p*-value from the test of group-specific correlation smooths. 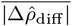 represents the average absolute difference between the predicted correlation curves of the two groups across evenly spaced effective niche values (*E*_*t*_). max 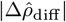 denotes the maximum absolute difference between the two predicted correlation trajectories across the effective-niche range. Larger values indicate stronger subtype-specific divergence in dynamic co-expression.

| | Gene 1 | Gene 2 | $p_{\text{adjust}}$ | $\max \vert \Delta\hat{\rho}_{\text{diff}}\vert $ | $\vert \Delta\hat{\rho}_{\text{diff}}\vert $ | | Gene 1 | Gene 2 | $p_{\text{adjust}}$ | $\max \vert \Delta\hat{\rho}_{\text{diff}}\vert $ | $\vert \Delta\hat{\rho}_{\text{diff}}\vert $ |
| --- | --- | --- | --- | --- | --- | --- | --- | --- | --- | --- | --- |
| 1 | APCDD1 | RND1 | < 0.001 | 0.67 | 0.30 | 25 | CNN3 | RBM3 | 0.005 | 1.00 | 0.37 |
| 2 | CFTR | EDN1 | < 0.001 | 0.67 | 0.32 | 26 | PFDN5 | RHOA | 0.005 | 0.36 | 0.15 |
| 3 | CFTR | FSTL3 | < 0.001 | 0.64 | 0.31 | 27 | CFTR | GADD45A | 0.005 | 0.53 | 0.25 |
| 4 | DUSP1 | TCIM | < 0.001 | 0.30 | 0.20 | 28 | LAMB3 | SERPING1 | 0.006 | 0.68 | 0.32 |
| 5 | MTRNR2L11 | NUPR1 | < 0.001 | 0.74 | 0.37 | 29 | CFB | CXCL6 | 0.006 | 0.27 | 0.12 |
| 6 | ACTN1 | ANXA3 | < 0.001 | 0.63 | 0.29 | 30 | CXCL2 | CXCL6 | 0.009 | 0.26 | 0.12 |
| 7 | CFB | FLNA | < 0.001 | 0.50 | 0.30 | 31 | ACTN1 | RHOA | 0.014 | 0.36 | 0.16 |
| 8 | CFB | FSTL3 | < 0.001 | 0.34 | 0.15 | 32 | CFB | SPIB | 0.014 | 0.48 | 0.21 |
| 9 | ANPEP | CFB | 0.001 | 0.40 | 0.17 | 33 | CXCR4 | EEF1B2 | 0.014 | 0.57 | 0.25 |
| 10 | C1R | CFB | 0.001 | 0.24 | 0.12 | 34 | MYH9 | NTN4 | 0.015 | 0.22 | 0.13 |
| 11 | CFB | PABPC1 | 0.001 | 0.21 | 0.10 | 35 | C1R | GATM | 0.015 | 0.58 | 0.30 |
| 12 | ANPEP | FSTL3 | 0.001 | 0.55 | 0.24 | 36 | CXCL2 | PFDN5 | 0.015 | 0.51 | 0.27 |
| 13 | ANXA3 | MYH9 | 0.001 | 0.56 | 0.29 | 37 | ACTN1 | FSTL1 | 0.020 | 0.83 | 0.32 |
| 14 | C1R | CXCL6 | 0.001 | 0.37 | 0.16 | 38 | DST | GPC1 | 0.020 | 0.46 | 0.20 |
| 15 | C5orf46 | CFB | 0.001 | 0.50 | 0.22 | 39 | ERBB2 | MORF4L1 | 0.020 | 0.42 | 0.18 |
| 16 | ACTN1 | FLNA | 0.001 | 0.55 | 0.27 | 40 | ASAH1 | RHOA | 0.026 | 0.33 | 0.14 |
| 17 | GATM | NOP53 | 0.001 | 0.56 | 0.25 | 41 | ATP5MD | CNN3 | 0.029 | 0.35 | 0.15 |
| 18 | BIRC3 | CFTR | 0.001 | 0.51 | 0.24 | 42 | GNAS | RBM3 | 0.030 | 0.31 | 0.13 |
| 19 | CXCL2 | FSTL3 | 0.002 | 0.31 | 0.14 | 43 | CFTR | LAMB3 | 0.036 | 0.43 | 0.18 |
| 20 | FLNA | FSTL1 | 0.003 | 0.46 | 0.21 | 44 | CFTR | SERPING1 | 0.036 | 0.42 | 0.22 |
| 21 | CFTR | DUSP1 | 0.003 | 0.38 | 0.27 | 45 | FSTL1 | GDF15 | 0.036 | 0.66 | 0.32 |
| 22 | CFB | HSP90B1 | 0.003 | 0.31 | 0.12 | 46 | ANPEP | SLC25A3 | 0.038 | 0.39 | 0.17 |
| 23 | DSP | EEF1B2 | 0.004 | 0.40 | 0.17 | 47 | C15orf48 | EEF1B2 | 0.038 | 0.55 | 0.27 |
| 24 | ACTN1 | EDN1 | 0.004 | 1.47 | 0.61 | 48 | GATM | GDF15 | 0.047 | 0.71 | 0.36 |

We additionally quantified the magnitude of the difference between the estimated correlation trajectories. Let 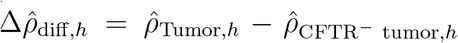, where 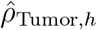 and 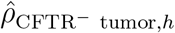 denote the predicted gene pair correlations at the *h*th effective-niche quantile value for tumor cells and CFTR^*−*^ tumor cells, respectively. The average absolute difference is

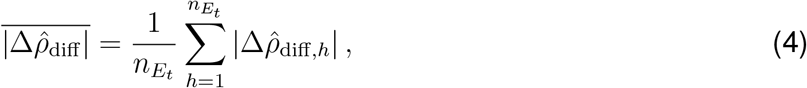

where 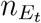 is the number of evenly spaced effective niche quantile values 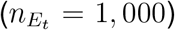 considered along the tumor-fibroblast effective niche, *E*_*t*_.

Because the distributions of *E*_*t*_ differed between tumor cells and CFTR^*−*^ tumor cells, we generated a common quantile grid of evenly spaced *E*_*t*_ values spanning the observed effective niche range. We then used the fitted NicheCorr model to predict correlation trajectories for both groups on this shared grid, enabling direct comparison of their dynamic co-expression patterns. In addition to the average difference, 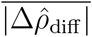, we computed max 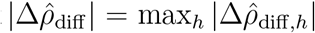, the maximum absolute difference between the two predicted correlation curves across all effective niche values. Because correlation values are bounded between −1 and 1, the theoretical upper bound of max 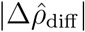is 2.

These results indicate that tumor-fibroblast spatial interactions reshape gene-gene coordination differently across tumor subtypes. Rather than simply altering individual gene expression levels, fibroblast proximity modulates the structure of gene co-expression networks in a subtype-specific manner.

To illustrate subtype-specific, niche-dependent co-expression patterns, we present the CFBFLNA gene pair as a representative example (Figure 8).

**Figure 8:**
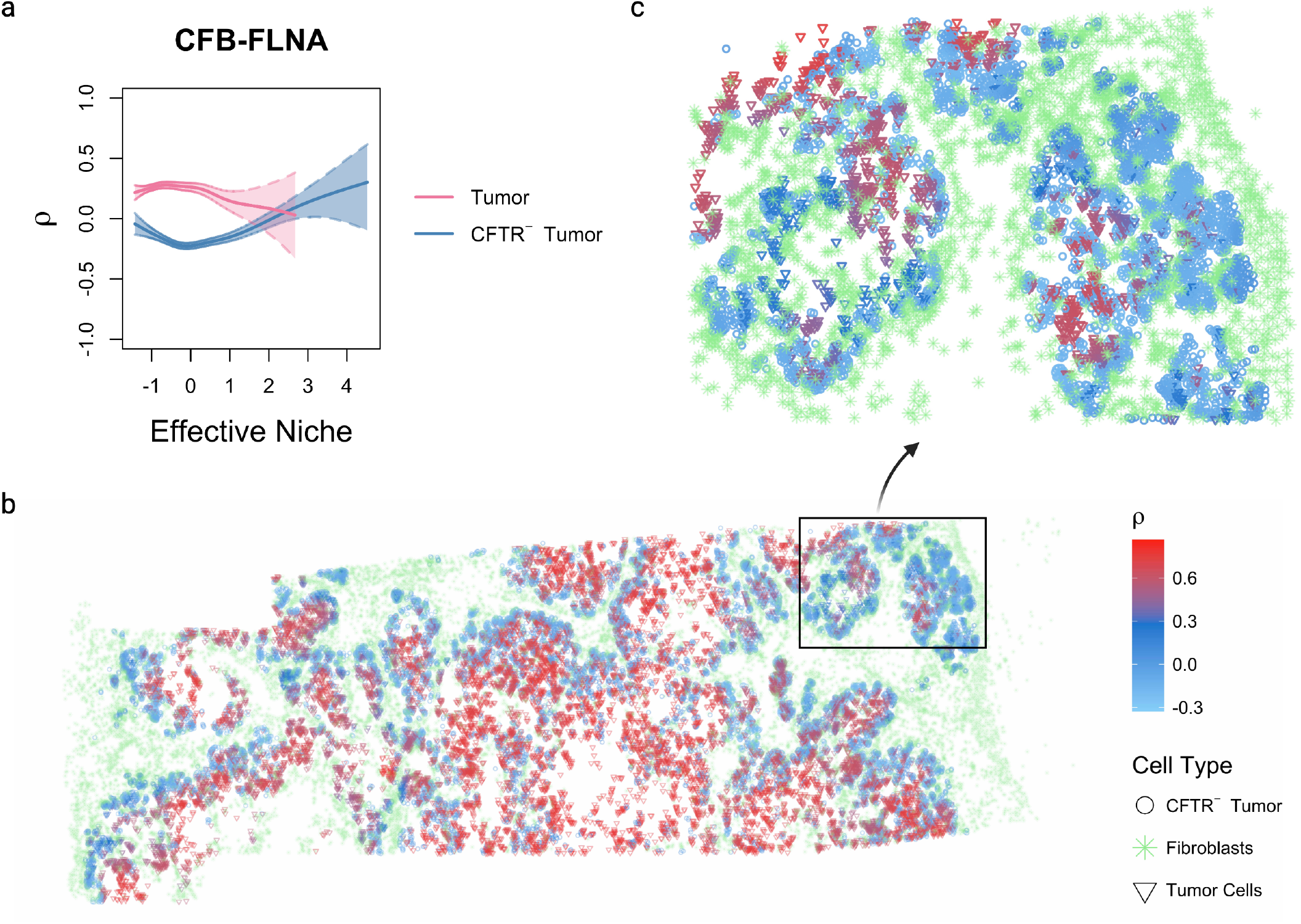
Subtype-specific, niche-dependent co-expression of CFB-FLNA. **a** Estimated correlation trajectories between CFB and FLNA across effective fibroblast niche strength (*E*_*t*_). Tumor cells are shown in red and CFTR^*−*^ tumor cells in blue. Solid lines represent the fitted NicheCorr correlation curves, and shaded regions denote 95% confidence intervals. **b** Spatial distribution of predicted CFB-FLNA correlation values mapped onto tissue coordinates. Cells are colored according to their locally predicted CFB-FLNA correlation, with symbols indicating cell identity. **c** Zoomed-in view of the representative tissue region highlighted in **b.** This regional view illustrates subtype-specific differences in CFB-FLNA co-expression within the same spatial microenvironment. Regions with higher effective fibroblast niche composition show increasing correlation in CFTR^*−*^ tumor cells, whereas tumor cells display stronger positive correlation in regions with lower effective fibroblast niche composition.

The correlation plot (Figure 8a) shows the estimated correlation trajectories across tumor-fibroblast niche strength, *E*_*t*_, for tumor cells (red) and CFTR^*−*^ tumor cells (blue). Shaded regions represent 95% confidence intervals. In tumor cells, CFB-FLNA exhibits a positive correlation at low effective fibroblast niche strength, corresponding to relatively few fibroblasts in the local neighborhood. This correlation gradually decreases as the effective fibroblast niche composition increases. In contrast, CFTR^*−*^ tumor cells display a different trajectory: the correlation is initially weakly negative and increases steadily with niche strength, eventually becoming positive at higher *E*_*t*_ values. These diverging trends indicate that fibroblast proximity modulates the coordination between CFB and FLNA differently across the two tumor subtypes.

The spatial plot (Figure 8b) projects the predicted correlation values onto tissue coordinates. Tumor cells are shown as triangles and CFTR^*−*^ tumor cells as circles, with color representing the locally predicted co-expression. Regions with higher effective fibroblast niche composition, defined as a greater distance-weighted abundance of fibroblasts in the local neighborhood, exhibit distinct spatial patterns between the two groups. Tumor cells tend to show positive CFB-FLNA correlation in areas with low effective fibroblast niche composition, whereas CFTR^*−*^ tumor cells display increasing correlation in regions with higher fibroblast niche composition.

The zoomed-in view (Figure 8c) highlights a representative tissue region in which these subtype-specific patterns are evident. Tumor cells in areas with relatively low fibroblast niche composition predominantly exhibit positive CFB-FLNA correlation (red). As the effective fibroblast niche composition increases, their correlation decreases (purple). In contrast, CFTR^*−*^ tumor cells show progressively stronger correlation as the local fibroblast niche composition increases (blue). This pattern illustrates how fibroblast proximity locally reshapes gene-gene coordination in a subtype-dependent manner.

This spatial visualization demonstrates that subtype-specific differences in co-expression are structured by the local microenvironment, indicating that variation in fibroblast niche composition systematically reshapes gene-gene coordination within the tissue architecture.

Together, these results show that fibroblast proximity reshapes gene-gene coordination in a subtype-dependent manner. Rather than uniformly altering gene expression levels, tumor-fibroblast interactions differentially alter the co-expression relationship between CFB and FLNA in tumor cells and CFTR^*−*^ tumor cells, highlighting microenvironment-driven transcriptional heterogeneity.

#### Ligand-receptor analysis

To identify LR interactions associated with subtype-specific co-expression changes captured by NicheCorr, we extended the Niche-LR framework to a subtype-aware setting. Rather than yielding a single chi-square statistic for each gene pair, the pancreatic cancer analysis estimates group-specific correlation smooths. The key signal is the extent to which each gene’s niche-dependent co-expression pattern with other genes differs between tumor cells and CFTR^*−*^ tumor cells. We therefore derived a differential gene-level statistic that quantifies this between-subtype divergence for each gene and used these statistics as input to the Niche-LR framework.

We visualized the significant gene pairs as an undirected network (Figure 9). Each node represents a gene, and each edge denotes a gene pair whose co-expression response to the fibroblast effective niche, *E*_*t*_, differs significantly between the two tumor subtypes. To highlight LR associations, nodes are displayed as pie charts, with each colored sector corresponding to a known LR interaction involving that gene. Candidate LR interactions identified using the subtype-aware Niche-LR framework, together with their top-ranked associated gene sets, are summarized in Table 5.

**Table 5:** Candidate LR interactions associated with subtype-specific, fibroblast niche-dependent co-expression patterns in pancreatic cancer. LR interactions were identified using the subtype-aware Niche-LR framework with differential gene-level statistics derived from NicheCorr. For each LR pair, the top gene set lists the highest-ranked genes associated with the interaction.

| Ligand | Receptor | top gene sets |
| --- | --- | --- |
| CD34 | SELL | DUSP1,CXCR4,GDF15,GADD45A,GNAS |
| EDN1 | EDNRB | DUSP1,EDN1,GDF15,GADD45A,ERBB2 |
| IL1B | IL1R2 | CFB,CXCL2,TCIM,CXCL6,BIRC3 |
| SERPING1 | SELE | CXCL2,EDN1,GDF15,GADD45A,CXCR4 |
| TFPI | F3 | DUSP1,GADD45A,BIRC3,C15orf48,ACTN1 |
| TNC | EGFR | DUSP1,EDN1,GDF15,GADD45A,CXCR4 |

**Figure 9:**
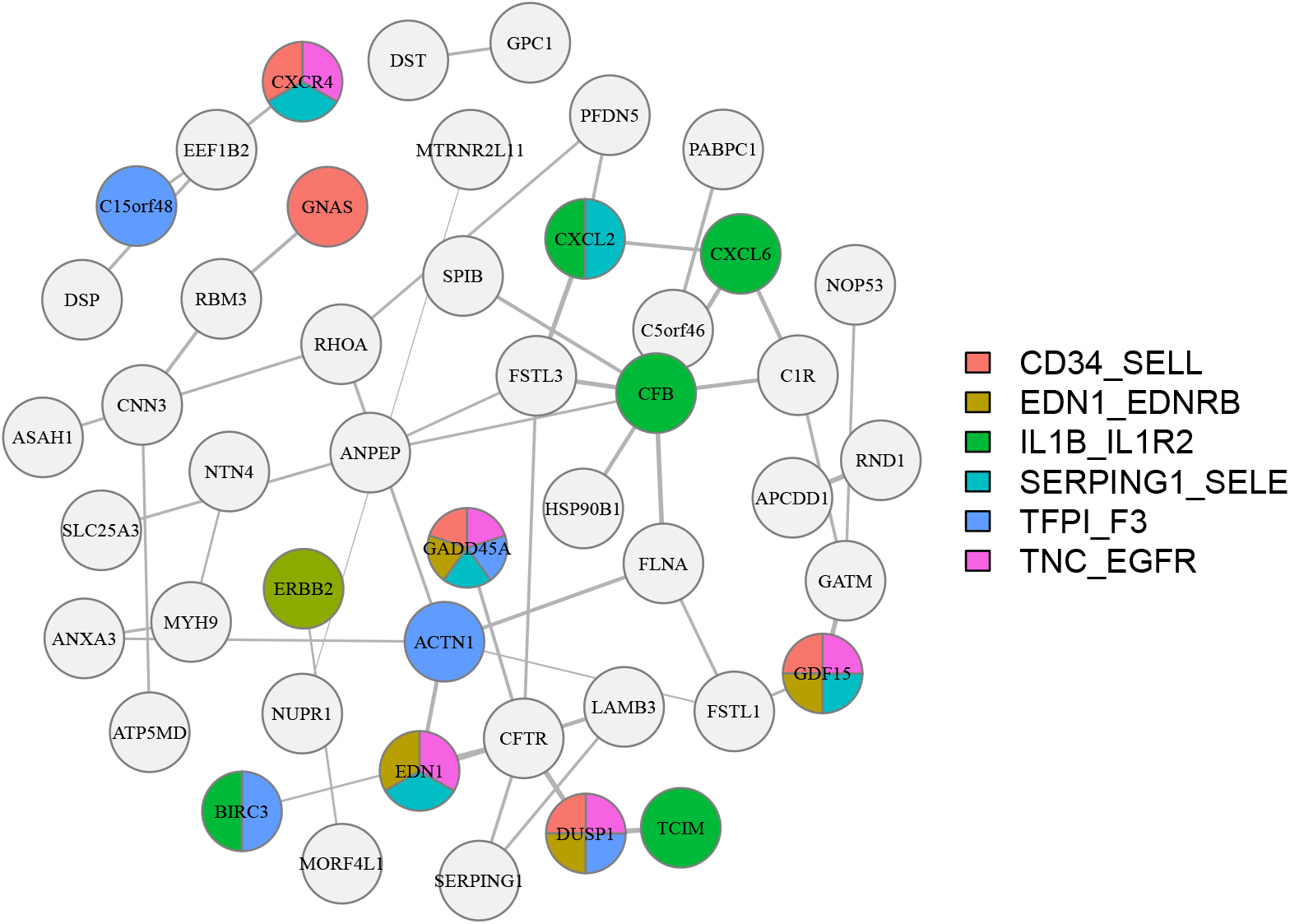
Subtype-differential gene network in the fibroblast effective niche. Nodes represent genes whose responses to *E*_*t*_ differ significantly between tumor cells and CFTR^*−*^ tumor cells. Edges connect gene pairs with subtype-differential niche responses, with edge width proportional to the magnitude of the differential statistic. Nodes are displayed as pie charts, with each colored sector corresponding to a known LR interaction involving that gene. This visualization highlights genes with potentially multifunctional signaling roles across the two tumor subtypes.

The network reveals hub genes with strong subtype-differential niche responses, including CFB, CFTR, and GDF15. These genes are centrally connected to multiple other genes, suggesting coordinated involvement in how tumor cells and CFTR^*−*^ tumor cells respond differently to their fibroblast microenvironment. Several of the identified LR pairs have established relevance in the pancreatic tumor microenvironment.

These results suggest that spatial proximity to fibroblasts reshapes gene co-expression programs differently in tumor cells and CFTR^*−*^ tumor cells. Genes involved in cell-cell communication show particularly strong subtype-specific changes, indicating that the two tumor subtypes may engage distinct LR signaling programs in response to their local fibroblast niche. This highlights the ability of NicheCorr to resolve subtype-level differences in microenvironment-driven co-expression patterns that may be obscured when tumor cells are treated as a single homogeneous population.

### Fibroblast gene expression as a covariate

Having identified 48 gene pairs with significantly different niche-dependent co-expression trajectories between tumor cells and CFTR^*−*^ tumor cells, we next investigated which fibroblast-derived genes in the local microenvironment were associated with these differential co-expression patterns.

To characterize the fibroblast transcriptional environment surrounding each tumor cell, we computed a spatially weighted average of fibroblast gene expression using a Gaussian kernel. For an index cell *i* and neighboring cell type *t* (fibroblasts in this analysis), the local expression of gene *g*_3_ among nearby cells of type *t* is summarized in Equation (6).

We applied an expression threshold of 0.1 and retained fibroblast genes expressed in at least 10% of cells in both the CFTR^*−*^ tumor and tumor cell populations. For each of the 48 significant gene pairs, we tested candidate fibroblast niche genes by modeling both the marginal distributions and the copula dependence parameter as functions of cell group, smooth nonlinear effects of the fibroblast niche gene, and group-specific smooth interactions. We performed analysis using the gene expression values in the fibroblast as a covariate in Equation (2)

To account for spatial autocorrelation, we additionally included MEMs as covariates in the marginal and dependence predictors. This formulation directly tests whether fibroblast-derived signals modulate the co-expression structure of each gene pair in a cell-type-specific manner, linking specific fibroblast genes to the differential co-expression patterns identified in the effective niche analysis.

Figure 10 summarizes the resulting third-gene modulation network, in which fibroblast-derived genes are connected to the tumor gene pairs whose co-expression they modulate. Nodes are displayed as pie charts colored by role: blue denotes tumor genes participating in subtype-differential co-expression pairs, and orange denotes fibroblast niche genes that modulate those pairs (third genes). Solid edges connect the two members of a significant tumor gene pair; dashed edges connect a fibroblast third gene to each member of the pair it modulates. Some genes act as both a tumor-pair member and a fibroblast modulator; these nodes are split blue and orange and carry both solid and dashed edges, reflecting their dual role in the network.

**Figure 10:**
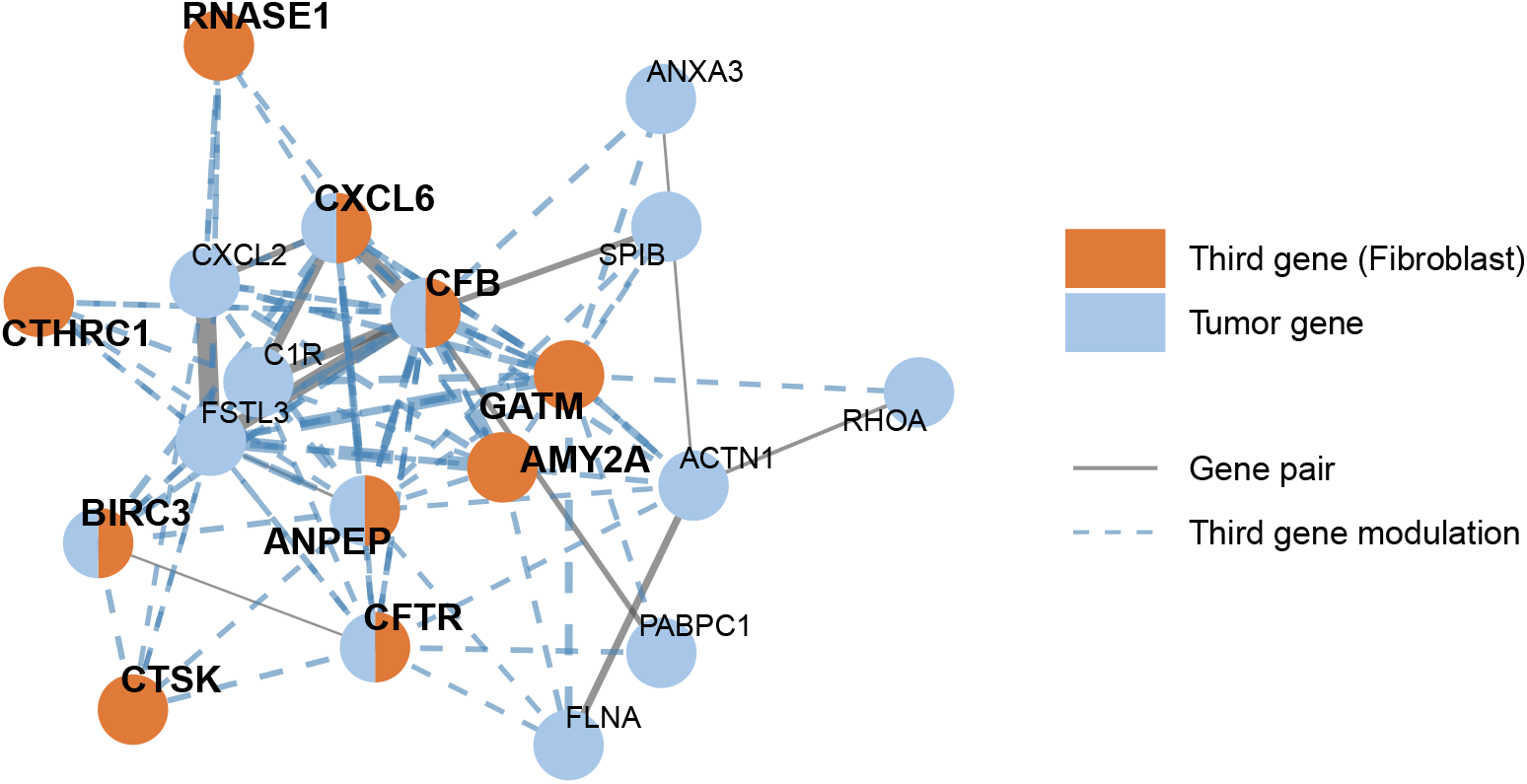
Third-gene modulation network in the fibroblast effective niche. Nodes are displayed as pie charts colored by role. Solid edges connect tumor gene pairs (blue nodes) with subtype-differential niche-dependent co-expression; edge width is proportional to the differential test statistic. Orange nodes represent fibroblast-derived third genes whose expression significantly modulates the co-expression of the connected gene pair differently between tumor and CFTR^*−*^ tumor cells. Dashed edges connect each third gene to the two members of the gene pair it modulates. Genes acting as both a tumor-pair member and a fibroblast modulator are split blue and orange, reflecting their dual role. Results are based on BH adjusted *p <* 0.05.

In total, 1681 gene pair–fibroblast gene triplets were significant at BH-adjusted *p <* 0.05, involving 199 distinct fibroblast genes. To highlight the strongest signals, Figure 10 and Table 3 (Supplementary Materials) show the triplets in the top 5% of test statistics among the ten fibroblast genes modulating the most gene pairs, giving 50 triplets spanning 13 tumor gene pairs. GATM was the most connected fibroblast modulator, associated with 11 of the 13 pairs, followed by AMY2A (7), ANPEP (6), CFTR (6), and CFB (5). Among tumor gene pairs, CFB–FSTL3 was modulated by the largest number of fibroblast genes (8), and it accounted for the two strongest triplets overall. For example, in the triplet CFB–FSTL3 with fibroblast AMY2A (*χ*^2^ = 106.89), the relationship between AMY2A expression in neighboring fibroblasts and the CFB–FSTL3 correlation differs significantly between tumor cells and CFTR^*−*^ tumor cells, indicating that the fibroblast niche modulates the co-regulation of this pair in a subtype-specific manner. Five genes (CFB, ANPEP, BIRC3, CFTR, and CXCL6) appeared both as members of differential tumor gene pairs and as fibroblast modulators, suggesting that they participate in tumor–fibroblast crosstalk in both directions.

## Discussion

We developed NicheCorr, a copula-based framework that models how gene–gene co-expression varies with spatial niche composition. Rather than focusing solely on mean expression differences, our method explicitly captures nonlinear, niche-dependent changes in gene–gene correlation. By integrating niche composition, copula-based dependence modeling, and smooth correlation trajectories, NicheCorr provides a principled way to quantify how microenvironmental proximity reshapes transcriptional coordination within cells.

A key strength of NicheCorr is its flexibility. The copula model separates marginal expression modeling from dependence modeling, allowing NB margins to accommodate overdispersed and zero-inflated count data while capturing complex, nonlinear correlation dynamics through smooth functions. This avoids restrictive linear assumptions and enables detection of subtle but biologically meaningful trajectory-dependent co-expression patterns.

The framework also provides direct interpretability. By estimating smooth correlation curves along the effective niche, the method yields intuitive visualizations of how gene–gene co-expression varies across the niche. This enables direct comparison between cell types or tumor subtypes, as demonstrated in the pancreatic cancer analysis, where tumor and CFTR^*−*^ tumor cells exhibited distinct niche-dependent correlation trajectories.

NicheCorr demonstrates strong statistical performance. Simulation studies show robust Type I error control and substantially higher power to detect dynamic correlation compared with existing approaches, particularly when mean expression varies nonlinearly with effective niche. The ability to model marginal mean trends explicitly prevents spurious detections driven by trajectory-dependent mean shifts.

The effective niche depends on the kernel bandwidth parameter, which determines the spatial scale of interaction. In practice, we recommend selecting the smallest bandwidth at which niche structure stabilizes, assessed using cosine distance between effective niche matrices across candidate bandwidths. This approach balances biological resolution and stability while avoiding over-smoothing. Density visualization of niche distributions can further ensure that meaningful heterogeneity is preserved.

Although demonstrated on CosMx and Xenium datasets, NicheCorr is broadly applicable to spatial transcriptomics at single-cell or spot-level resolution. The framework naturally extends to spot-level data by treating spots as observational units and computing the effective niche using spatial neighborhood weights. Additionally, it can accommodate multiple niche covariates simultaneously (e.g., immune abundance, fibroblast density, hypoxia scores), enabling modeling of more complex microenvironmental interactions.

NicheCorr enables multiple downstream strategies: ranking gene pairs by the significance of dynamic correlation change; constructing niche-dependent gene co-expression networks; and integrating with ligand–receptor inference frameworks.

The method is computationally feasible for moderate gene sets. All simulations and data analyses were conducted on a high-performance computing cluster in parallel, with each node configured with a single CPU. On average, model fitting took 1.96 minutes per gene pair for 4,500 observations. For large-scale studies, we recommend gene pre-filtering (e.g., based on expression level or biological relevance) to reduce computational burden while preserving signal. In practice, NicheCorr can be efficiently applied to whole-genome scRNA-seq datasets either through pathway-guided analysis of biologically relevant gene sets or via genome-wide pairwise screening using pre-filtering strategies such as Liquid Association Coefficient (LAC)^12^ or fastLA^13^ to reduce computational burden while retaining dynamic co-expression signals.

Adjacent cells share overlapping neighborhoods, so their niche values are spatially autocorrelated by construction. To address this, we included MEMs as spatial covariates in the marginal and dependence predictors. MEMs were derived from the eigenvectors of a spatial weights matrix built on a symmetric *k*-nearest-neighbor graph of the cell coordinates. We retained the leading eigenvectors, which capture the broad-scale spatial structure at the scale on which the niche is smoothed, removing global autocorrelation while preserving local variation. The same fixed set was applied to every gene pair. We recommend mapping the retained eigenvectors back onto the tissue coordinates to confirm that they represent broad-scale structure; the number retained depends on the spatial organization of the tissue and is therefore chosen per dataset.

## Methods

### Overview of the NicheCorr Framework

We developed NicheCorr, a copula-based statistical framework for characterizing how genegene co-expression relationships vary continuously with the spatial microenvironment. The central idea is to treat the pairwise correlation between two genes as a smooth function of a cell-level covariate **x**_*i*_ that summarizes the microenvironment surrounding cell *i*, so that coexpression is allowed to change across the tissue rather than being assumed constant. The framework accommodates any such covariate; in this work, we use two: the spatial niche composition (Section Spatial Niche Quantification) and the spatially weighted average expression of a neighboring-cell gene (Section Third-Gene Modulation of Co-expression).

The framework integrates three core components: (1) quantification of the microenvironment covariate **x**_*i*_; (2) joint distributional modeling of gene-pair expression through a Gaussian copula with NB marginal distributions, which separates each gene’s marginal expression from the dependence between genes; and (3) smooth, nonparametric modeling of the copula dependence parameter as a function of **x**_*i*_, including group-specific effects that allow the co-expression trajectory to differ across cell subtypes. Modeling the dependence parameter directly is what lets NicheCorr detect changes in co-expression that are not driven by shifts in mean expression. Downstream outputs include niche-dependent co-expression networks and integration with LR inference.

### Spatial Niche Quantification

We first take the microenvironment covariate **x**_*i*_ to be the effective niche composition surrounding each cell, adopting the niche construction approach from Niche-DE^1^. Assume that we have *N* cells and *G* genes. For each cell *i*, where *i* = 1, … , *N* , let **r**_*i*_ ∈ℝ^2^ denote its spatial coordinates and let *T*_*i*_ ∈{1, … , *T}* denote its cell-type label. The effective niche abundance of cell type *t* surrounding cell *i* is computed as

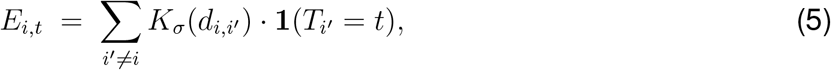

where 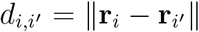 is the Euclidean distance between cells *i* and *i*^*′*^, and *K*_*σ*_(·) is a Gaussian kernel with bandwidth *σ*. Thus, *E*_*i,t*_ can be interpreted as the distance-weighted abundance of cell type *t* among cells surrounding cell *i*. Across all *N* cells and *T* cell types, this yields an effective niche matrix **E** ∈ℝ^*N×T*^ , with **x**_*i*_ = (*E*_*i*,1_, … , *E*_*i,T*_)^*⊤*^. Each entry quantifies the weighted local abundance of a given cell type around an index cell, with spatially proximate cells contributing more than distant cells. Following Niche-DE^1^, each column of **E** is then standardized across cells (centered to mean zero and scaled to unit variance) before model fitting. Although the raw kernel-weighted abundance in Equation (5) is nonnegative, the standardized values can therefore be negative, with negative values indicating a below-average abundance of cell type *t* around cell *i*. All values of *E*_*i,t*_ (and *E*_*t*_) reported in this paper are on this standardized scale.

### Third-Gene Modulation of Co-expression

Beyond the effective niche composition, we consider a second **x**_*i*_: the spatially weighted average expression of a gene in neighboring cells of a given type. This allows us to ask whether the coexpression of a gene pair (*g*_1_, *g*_2_) in an index cell is modulated by the local expression of a third gene *g*_3_ in a surrounding cell type *t* (e.g., fibroblasts).

For an index cell *i* and neighboring cell type *t*, the local expression of gene *g*_3_ among nearby cells of type *t* is summarized as

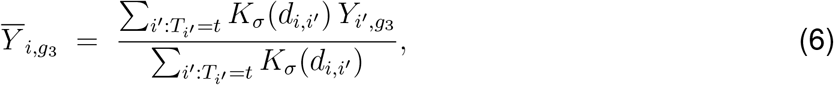

where 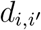 is the Euclidean distance between cells *i* and *i*^*′*^, *K*_*σ*_(·) is a Gaussian kernel with bandwidth *σ*, and 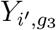 is the expression count of gene *g*_3_ in cell *i*^*′*^. The weights are normalized to sum to one for each index cell, so that 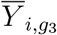 represents the local, spatially weighted expression of gene *g*_3_ in the type-*t* microenvironment surrounding cell *i*.

### Bandwidth Selection for Effective Niche Estimation

The effective niche and the spatially weighted average expression of a neighboring-cell gene were both computed using a kernel-based method that weights nearby cells according to their spatial distance, with the kernel bandwidth controlling the spatial scale over which cells are aggregated. Because the choice of bandwidth directly affects the inferred spatial structure, we evaluated multiple candidate bandwidths and applied the same selection procedure to both covariates.

For each bandwidth, a cell-level matrix was computed using a Gaussian kernel, quantifying the weighted contribution of neighboring cells to the local neighborhood of every cell. To compare these matrices across bandwidths, we flattened each into a vector and computed pairwise cosine distances, which emphasize differences in the overall spatial pattern rather than absolute magnitude.

Bandwidth selection was then guided by two criteria. First, we compared the matrix at each bandwidth to that obtained at the largest bandwidth and selected the smallest bandwidth at which the cosine distance dropped below a predefined threshold (0.1), an elbow point beyond which further increases yielded no meaningful change in spatial structure. Second, we examined the distribution of the resulting covariate across bandwidths using density plots to confirm that the selected bandwidth preserved biologically meaningful variation while avoiding over-smoothing.

### Joint Modeling Framework for Gene Co-expression

To characterize microenvironment-specific co-expression, we modeled the joint distribution of gene pairs. For a gene pair (*g*_1_, *g*_2_), let 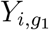 and 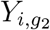 denote the observed counts for genes *g*_1_ and *g*_2_ in cell *i*, respectively. We used a Gaussian copula to capture the dependence structure between gene pairs, with NB marginal distributions to account for overdispersed count data common in spatial transcriptomics:

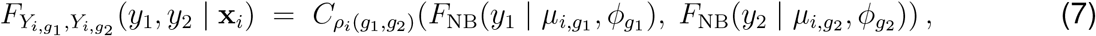

where *C*_*ρ*_(·, ·) is a Gaussian copula with correlation parameter *ρ*, and *F*_NB_(·|*µ, ϕ*) is the cumulative distribution function of an NB distribution with mean *µ* and overdispersion parameter *ϕ*. Here **x**_*i*_ denotes the vector of microenvironment covariates for cell *i*. In this work each fit uses a single covariate, which we write as the scalar *x*_*i*_: in the niche analysis *x*_*i*_ = *E*_*i,t*_ for a chosen niche cell type *t*, and in the third-gene analysis 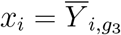. The NB marginals accommodate overdispersed count data and can model the many zero counts commonly observed in spatial transcriptomics without requiring a separate zero-inflation component.

The copula structure separates marginal behavior from the dependence structure, allowing gene-gene correlations to vary flexibly with the microenvironment while each marginal mean is independently modeled as a smooth function of the same covariate. This separation ensures that detected co-expression changes are not confounded by shifts in marginal expression levels. NicheCorr models the copula dependence parameter as a smooth function of the microenvironment covariate. To account for residual spatial autocorrelation between cells, we additionally include a set of MEMs as linear covariates. MEMs are eigenvectors of a spatial weights matrix derived from a nearest-neighbor graph of the cells; the retained leading eigenvectors remove global autocorrelation while preserving the local, covariate-driven variation of interest. The same fixed set of MEMs is used across all gene pairs within an analysis, and enters the predictors as linear terms 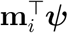, where **m**_*i*_ is the vector of MEM values for cell *i* and ***ψ*** is the corresponding coefficient vector.

Each gene pair is fitted separately, so for readability we suppress the gene-pair index on the model parameters below; every intercept, smooth function, and coefficient vector with a *ρ* subscript is specific to the pair (*g*_1_, *g*_2_). For a gene pair (*g*_1_, *g*_2_), we specify the additive predictor for the dependence parameter, the single-covariate form of Equation (1) with MEM adjustment, as

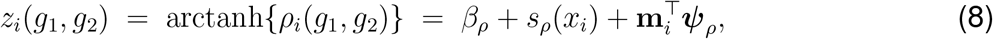

where *β*_*ρ*_ is a gene-pair-specific intercept (written *β*_0_ in Equation (1)), *s*_*ρ*_(·) is a thin-plate regression spline that flexibly captures the nonlinear dependence of correlation on the covariate, and ***ψ***_*ρ*_ are the corresponding MEM coefficients. Other spline bases, such as cubic regression splines, B-splines, and P-splines, can also be used. The correlation is recovered through the inverse link,

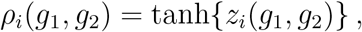

which ensures that *ρ*_*i*_(*g*_1_, *g*_2_) ∈ (−1, 1).

Similarly, the marginal mean expression of each gene is modeled as

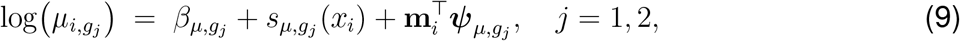

where 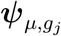 are the MEM coefficients for the marginal mean of gene *g*_*j*_. The number of MEMs is not fixed across analyses; it is chosen per dataset.

For each gene pair (*g*_1_, *g*_2_), the full parameter vector is

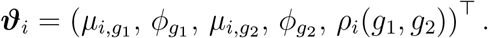

The parameters are estimated jointly by maximizing the penalized log-likelihood,

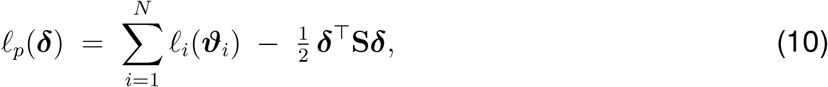

where *l*_*i*_(·) is the log-likelihood contribution of cell *i* under Equation (7), ***δ*** collects all spline basis coefficients for gene pair (*g*_1_, *g*_2_), and **S** is a block-diagonal smoothing penalty matrix with smoothing parameters selected by generalized cross-validation^14–16^.

### Multi-group Extension for Subtype Comparison

To compare co-expression across cell subtypes (e.g., tumor cells versus CFTR^*−*^ tumor cells), we extended the model to include group-specific smooth effects. Let *γ*_*i*_ ∈{1, … , *Q}* denote the subtype label of cell *i*, where *Q* is the total number of groups and group 1 serves as the reference group.

For a gene pair (*g*_1_, *g*_2_), let *g*_*j*_ denote gene *j*, where *j* ∈ {1, 2} . The additive predictor for the marginal mean expression of gene *g*_*j*_ becomes

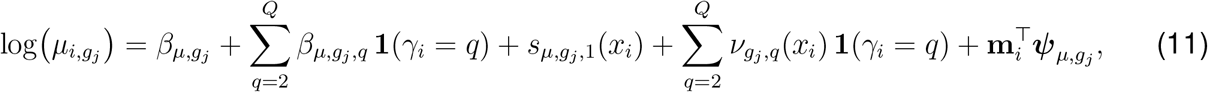

where 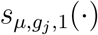 is the smooth mean trajectory for gene *g*_*j*_ in the reference group, 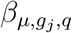 captures the intercept shift for group *q*, and 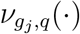 is the deviation smooth for group *q* relative to the reference group. Thus, the group-specific smooth component for gene *g*_*j*_ in group *q* is 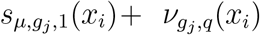.

The additive predictor for the copula correlation parameter becomes

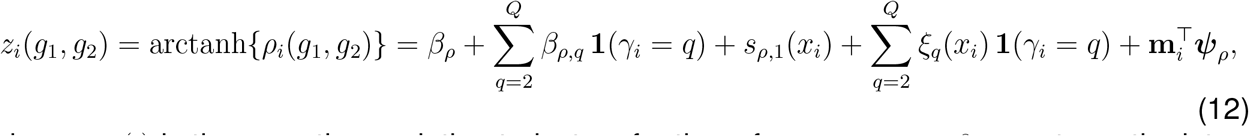

where *s*_*ρ*,1_(·) is the smooth correlation trajectory for the reference group, *β*_*ρ,q*_ captures the intercept shift for group *q* relative to the reference group, *ξ*_*q*_(·) is a group-specific deviation smooth that models how the correlation trajectory of group *q* departs from the reference trajectory, and 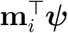 is the MEM adjustment. As above, all of these *ρ*-model terms are specific to the gene pair (*g*_1_, *g*_2_).

Thus, the group-specific smooth component for the correlation in group *q* is *s*_*ρ*,1_(*x*_*i*_) + *ξ*_*q*_(*x*_*i*_). This parameterization has a direct inferential advantage: each deviation smooth, *ξ*_*q*_(·) and 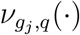, is a penalized spline term whose significance can be assessed using a chi-square test based on its associated effective degrees of freedom. A significant *ξ*_*q*_(·) indicates that group *q* has a statistically distinct covariate-dependent correlation trajectory from the reference group, whereas a significant 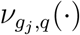 indicates differential covariate-dependent mean expression dynamics for gene *g*_*j*_.

### Hypothesis Testing and Multiple Testing Correction

For each gene pair (*g*_1_, *g*_2_), we tested whether its correlation varied with the covariate:

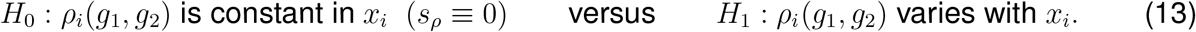

The resulting *p*-values were adjusted across all tested gene pairs using the BH procedure to control the false discovery rate (FDR), with an FDR threshold of 0.05.

In the multi-group setting, the parameterization in (12) provides a direct test for differential co-expression between subtypes. Each group-specific deviation smooth *ξ*_*q*_(·) is assessed by a chi-square test on its effective degrees of freedom, and a significant term indicates that group *q* has a correlation trajectory that differs from the reference group. As in the single-group case, the resulting *p*-values were adjusted across gene pairs using the BH procedure at an FDR threshold of 0.05.

### Niche-Dependent Co-expression Networks and Ligand-Receptor Analysis

Significant gene pairs were assembled into an undirected co-expression network, in which nodes represent genes and edge weights are proportional to the chi-square test statistic, reflecting the strength of evidence for covariate-varying correlation. Hub genes, defined as genes appearing in multiple significant pairs, were identified as candidates for coordinated microenvironmental regulation.

To link detected co-expression changes to upstream signaling, we extended the Niche-LR framework^1^ to use NicheCorr chi-square statistics in place of Niche-DE *t*-statistics. For each gene *g* involved in the significant pairs, we computed a gene-level statistic by summing the chisquare statistics across all significant pairs involving *g*:

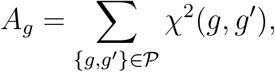

where *P* is the set of significant gene pairs, so that genes participating in more significant pairs receive larger aggregate statistics.

These gene-level statistics were then used as input for ligand-receptor (LR) enrichment scoring against NicheNet ligand-target matrices, identifying putative LR pairs whose downstream target genes showed the strongest covariate-dependent co-expression signals.

## RESOURCE AVAILABILITY

### Lead contact

Requests for further information and resources should be directed to and will be fulfilled by the lead contact, Shuyi Yang.

## Data and code availability

This paper analyzes existing, publicly available data. The CosMx SMI Human Liver FFPE dataset is available from NanoString^10^, and the Xenium Human Multi-Tissue and Cancer Panel pancreatic cancer dataset is available from 10x Genomics^11^.

