## Supplementary for "NicheCorr: Identifying Niche-Specific Differential Gene Co-Expression in Spatial Transcriptomics"

### NicheCorr Supplementary Appendix

## S1.

### S1.2

Table S1

Table S2

Figure S1

### S2. Binned Pearson Permutation Test for Non-Constant Correlation

#### Test Statistic

Let $\mathbf{y}_1 = (y_{11}, \dots, y_{1N})$ and $\mathbf{y}_2 = (y_{21}, \dots, y_{2N})$ denote the observed expression counts for two genes across $N$ cells, and let $E_{i,t}$ be the effective niche covariate of cell $i$ (written $E_t$ when the cell index is dropped). We test the null hypothesis $H_0$ that the Pearson correlation between the two genes is constant across all values of $E_t$, against the alternative that it varies with $E_t$.

To construct the test statistic, cells are partitioned into $L$ equally spaced bins along the $E_t$ axis. Within each non-empty bin $l$, the Pearson correlation $r_l$ is estimated from the cells assigned to that bin. To stabilise variance and normalise the sampling distribution of the correlation estimate, we apply the Fisher $z$-transformation:

$$z_l = \text{atanh}(r_l) = \frac{1}{2} \log\left(\frac{1+r_l}{1-r_l}\right), \quad l = 1, \dots, L. \quad (1)$$

Under $H_0$, the bin-level values $z_l$ fluctuate around a common mean. The observed test statistic summarises their variability:

$$T_{\text{obs}} = \sum_{l=1}^L (z_l - \bar{z})^2, \quad (2)$$

where $\bar{z} = L^{-1} \sum_l z_l$ is the cross-bin mean. A large $T_{\text{obs}}$ indicates that the correlation changes systematically across the $E_t$ axis, providing evidence against $H_0$.

#### Permutation Null Distribution

The null distribution of $T_{\text{obs}}$ is obtained by permutation. Under $H_0$, the correlation between the two genes does not depend on $E_t$, so the $E_t$ values are exchangeable among cells. For each permutation replicate $b = 1, \dots, B$, we draw a uniform random permutation of the $E_t$ values, reassign them to cells, re-bin, and recompute the test statistic $T_b^*$ via equations (1)–(2). This preserves the joint distribution of $(y_1, y_2)$ while destroying any dependence between the correlation level and $E_t$. The permutation $p$-value is:

$$p = \frac{1 + \sum_{b=1}^B \mathbf{1}(T_b^* > T_{\text{obs}})}{1 + B}, \quad (3)$$

where the $+1$ in both numerator and denominator incorporates the observed dataset itself as the identity permutation, ensuring that $p$ is a valid probability bounded away from zero<sup>1</sup>. We use $B = 1000$ permutations throughout, so that $p$-values lie on the grid $\{1/1000, 2/1000, \dots\}$ and the nominal $\alpha = 0.05$ threshold is exactly achievable.

### Selection of the Number of Bins

The number of bins $L$ governs a fundamental bias–variance trade-off. Too few bins reduce resolution and sacrifice power to detect localised changes in correlation; too many bins produce unstable within-bin estimates when individual bins contain few cells, inflating the type I error rate. We select $L$ through a simulation-based calibration procedure that is specific to the permutation test and the negative-binomial data structure. The goal is a single universal $L$ that is valid across all analyses without per-dataset customisation.

The calibration proceeds in two stages over a candidate grid $L \in \{5, 10, 15, 20, 25, 30, 35, 40, 45, 50\}$. In **Stage 1**, we simulate datasets under the null (constant correlation, $\lambda = 0$) and apply the test at nominal level $\alpha = 0.05$. Candidate values of $L$ for which the empirical rejection rate exceeds 0.05 are discarded. In **Stage 2**, we simulate datasets under a range of alternatives and record the empirical power for each $L$ that passed Stage 1. The optimal $L^*$ is the value that maximises average power across the alternatives:

$$L^* = \arg \max_L \overline{\text{power}}(L), \quad \text{subject to } \widehat{\text{typeI}}(L) \leq 0.05, \quad (4)$$

where $\overline{\text{power}}(L)$ denotes the average empirical power across the simulated alternatives. As a supplementary diagnostic, we verify that the null $p$-values are approximately uniform for each surviving $L$ using the Kolmogorov–Smirnov test. A non-uniform null $p$-value distribution under $H_0$ would signal that the permutation null is not adequately approximating the true null distribution for that bin count, regardless of the rejection rate criterion.

This procedure yields a $L^* = 30$ that is calibrated to the actual sampling variability of the test under the study design, and is stable across a wide range of effect sizes. The calibration is run once using the reference simulation model; the resulting $L^*$ is then applied uniformly to all datasets without further tuning.

### S3. Spatial Domain Biological Annotation

To biologically interpret the spatial domains identified by Leiden clustering, we assigned each domain a biological label based on cell type signature scoring. For each major cell type present in the dataset, we derived a set of marker genes by performing a Wilcoxon rank-sum test comparing each cell type against all others using the annotated cell type labels provided with the Xenium dataset. The top 30 marker genes per cell type were retained as the signature (Figure 2).

Cell type signatures were derived for the following populations: Tumor Cells, CFTR<sup>−</sup> Tumor Cells, Fibroblasts, Acinar Cells, Ductal Cells, Macrophages, T Cells, B Cells, Endocrine Cells

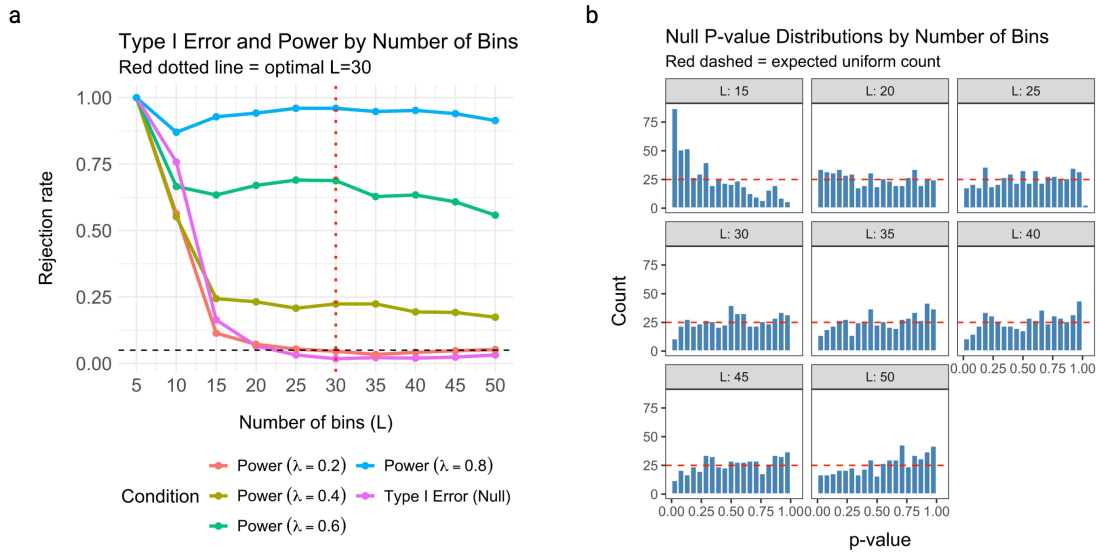

Figure 1: **a** Empirical rejection rates at nominal level $\alpha = 0.05$ as a function of the number of bins $L$, estimated from 500 simulation replicates. Pink: type I error under the null ($\lambda = 0$); other colors: power under four alternatives ($\lambda \in \{0.2, 0.4, 0.6, 0.8\}$). The horizontal dashed line marks $\alpha = 0.05$. The red dotted vertical line indicates the selected optimal $L^* = 30$, defined as the smallest $L$ at which the type I error is controlled at the nominal level, beyond which power is maximised. **b** Null $p$-value distributions for $L \in \{15, 20, 25, 30, 35, 40, 45, 50\}$. The red dashed horizontal line indicates the expected uniform count. Distributions become increasingly uniform as $L$ increases beyond 25, confirming that the permutation null is well-calibrated at $L^* = 30$.

(combining Endocrine 1 and Endocrine 2), and Metaplastic Cells. For each signature, we computed a per-cell score by calculating the average expression of the signature genes relative to a randomly sampled background gene set of comparable expression level, following the approach of Tirosh et al.<sup>2</sup>. Per-domain scores were obtained by averaging per-cell scores across all cells within each domain.

Each domain was assigned a biological label based on the highest mean signature score (Table 1, Table 2). Domains with a clearly dominant score were assigned a specific biological label, while domains with uniformly weak scores across all signatures were labeled as mixed. The resulting domain annotations are summarized in Table 2. For downstream NicheCorr analysis, we selected domains enriched in Tumor Cells (Domain 5), CFTR<sup>-</sup> Tumor Cells (Domain 8), and Fibroblasts (Domain 3), as these domains showed the strongest and most interpretable biological signals and were directly relevant to our analysis of fibroblast niche effects on tumor cell gene co-expression.

Table 1: Mean cell type signature scores per spatial domain. Scores reflect the average expression of cell type marker genes relative to a background gene set. Higher positive values indicate stronger enrichment of the corresponding cell type in that domain.

| Domain | Tumor | CFTR<sup>-</sup> Tumor | Stroma | Acinar | Ductal | Immune | Endocrine | Metaplastic |
| --- | --- | --- | --- | --- | --- | --- | --- | --- |
| 0 | 0.03 | 0.08 | 0.04 | 0.12 | 0.03 | 0.08 | 0.01 | 0.04 |
| 1 | 0.09 | 0.22 | -0.02 | 0.31 | 0.10 | 0.31 | 0.00 | 0.11 |
| 2 | 0.11 | 0.47 | -0.37 | 2.23 | 0.08 | -0.31 | 0.60 | 2.06 |
| 3 | 0.20 | 0.16 | 1.00 | 0.19 | 0.13 | 0.07 | 0.00 | -0.19 |
| 4 | 0.42 | 1.30 | -0.53 | 2.25 | 0.23 | -0.40 | 0.21 | 2.34 |
| 5 | 1.75 | 1.66 | -0.34 | 0.73 | 1.17 | -0.30 | 0.21 | 0.44 |
| 6 | 0.37 | 0.67 | -0.19 | 0.73 | 0.22 | -0.14 | 0.08 | 0.75 |
| 7 | 0.48 | 0.57 | -0.36 | 0.34 | 0.15 | -0.24 | 1.29 | 0.06 |
| 8 | 1.51 | 2.33 | -0.38 | 0.86 | 0.80 | -0.39 | 0.13 | 1.07 |
| 9 | 0.26 | 0.37 | 0.03 | 0.25 | 0.17 | -0.13 | 0.06 | 0.04 |
| 10 | 1.61 | 1.01 | -0.37 | 0.55 | 1.93 | -0.14 | 0.30 | 0.23 |

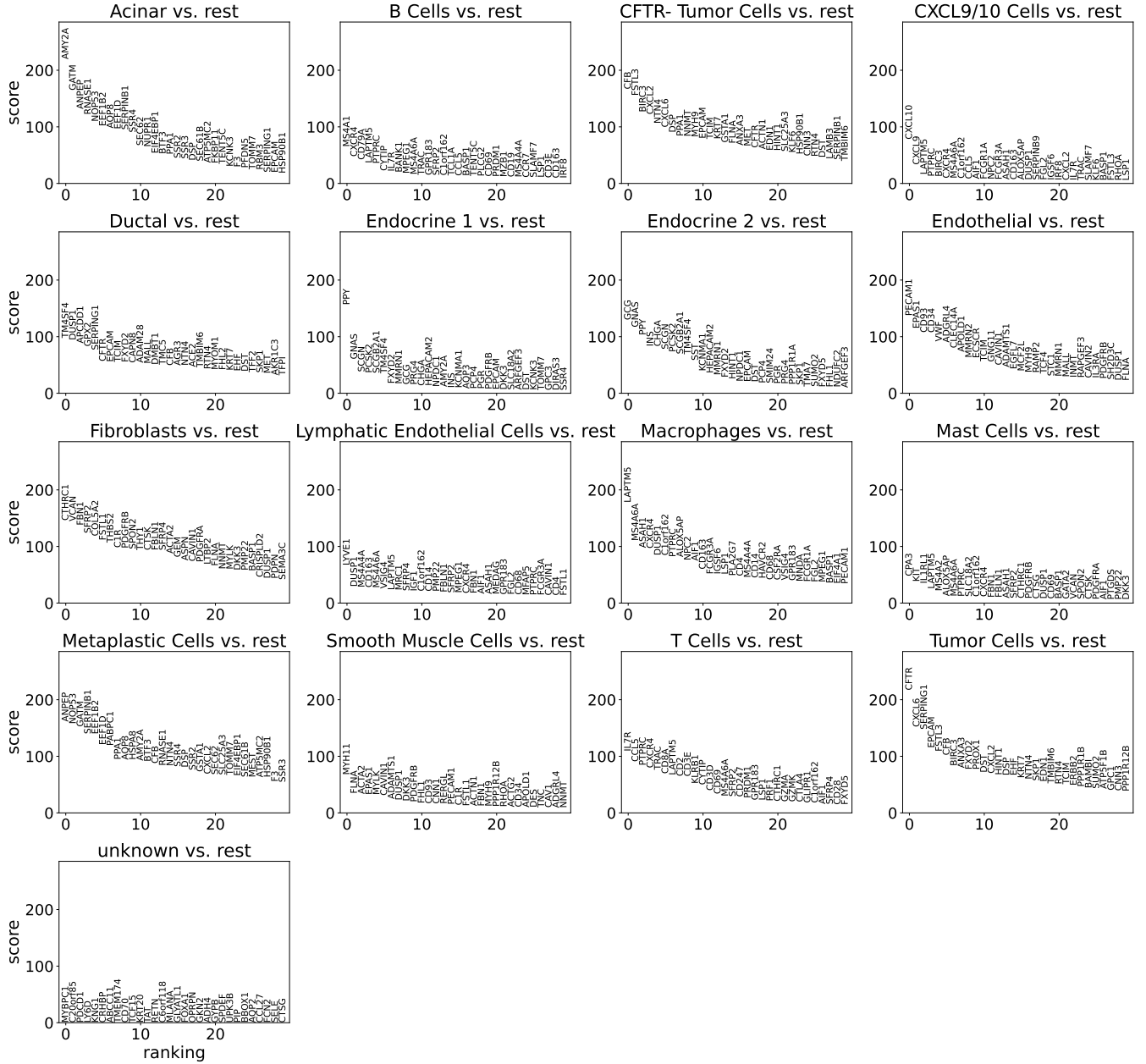

Figure 2: Top 30 marker genes per annotated cell type identified by Wilcoxon rank-sum test comparing each cell type against all others. The y-axis shows the Wilcoxon score and the x-axis shows gene ranking. Marker genes from these signatures were used to score spatial domains and assign biological labels.

Table 2: Biological labels assigned to spatial domains based on the highest mean cell type signature score. Domains with uniformly weak scores across all signatures are labeled as mixed.

| Domain | Highest Score | Value | Biological Label |
| --- | --- | --- | --- |
| 0 | acinar (0.12) | very weak | Mixed |
| 1 | immune (0.31) | weak | Mixed/immune |
| 2 | metaplastic (2.06) | very strong | Metaplastic niche |
| 3 | stroma (1.00) | strong | Stromal/fibroblast niche |
| 4 | metaplastic (2.34) | very strong | Metaplastic niche |
| 5 | tumor (1.75) | strong | Tumor-enriched niche |
| 6 | metaplastic (0.75) | moderate | Metaplastic/tumor mixed |
| 7 | endocrine (1.29) | strong | Endocrine/islet niche |
| 8 | CFTR<sup>-</sup> tumor (2.33) | very strong | CFTR<sup>-</sup> tumor niche |
| 9 | CFTR<sup>-</sup> tumor (0.37) | weak | Mixed/tumor |
| 10 | ductal (1.93) | strong | ductal-enriched niche |

### S4. Additional Results

#### 0.1 Pancreatic Cancer Xenium Dataset Analysis

Table 3 lists the gene pair–fibroblast gene triplets shown in the third-gene modulation network. Each row corresponds to a tumor gene pair ($g_1, g_2$) whose niche-dependent co-expression is significantly modulated by the expression of a fibroblast gene ($g_3$) in the local microenvironment, with the modulation differing between tumor and CFTR<sup>-</sup> tumor cells. GATM was the most frequent fibroblast modulator (11 triplets), and CFB–FSTL3 was the most frequently modulated tumor gene pair (8 triplets), including the two strongest associations, with AMY2A ($\chi^2 = 106.89$) and GATM ($\chi^2 = 106.62$).

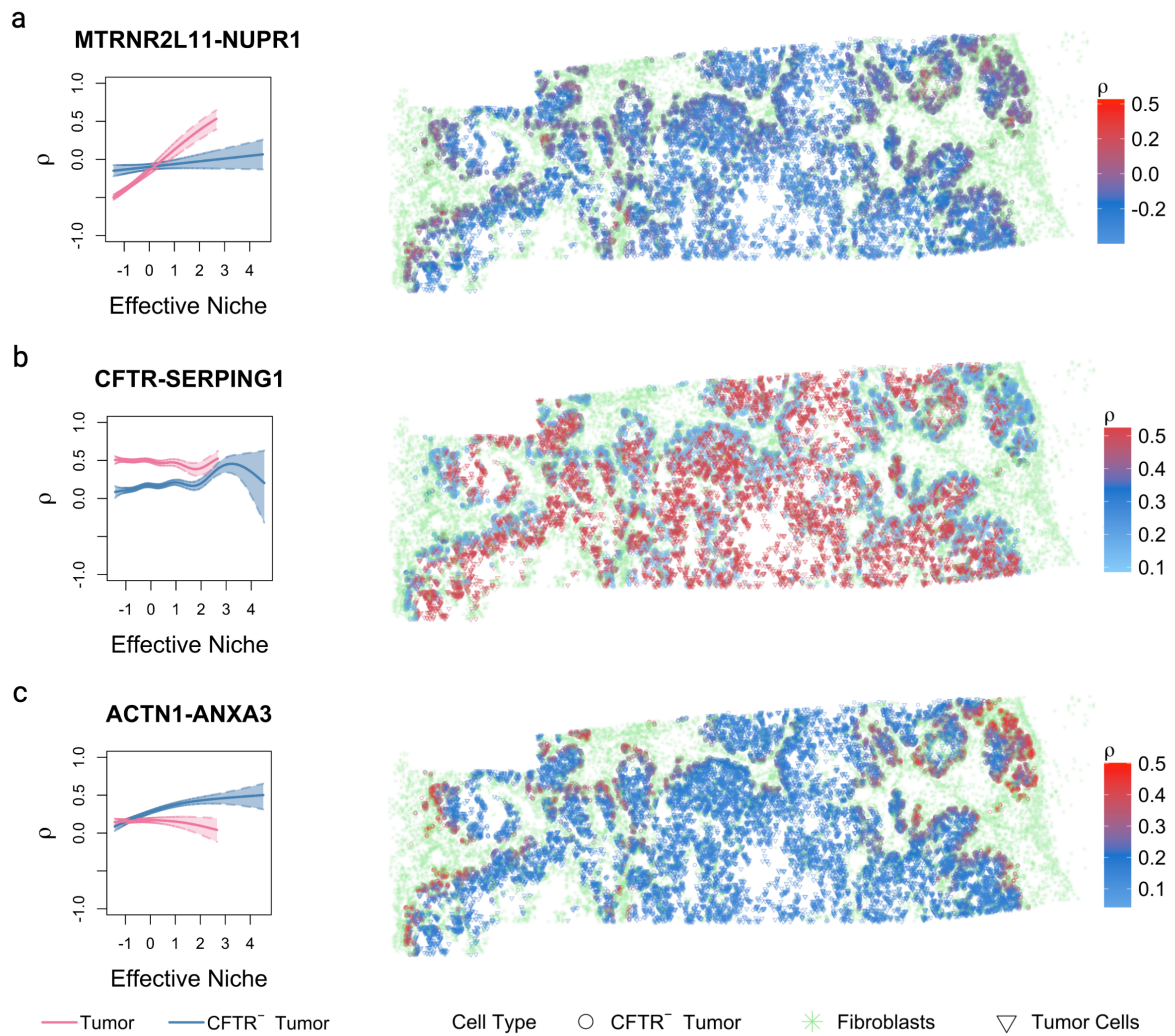

Figure 3:

Table 3: Significant third-gene modulation triplets (BH-adjusted $p < 0.05$), restricted to the top 5% of test statistics and the ten most connected fibroblast genes. $g_1, g_2$: tumor gene pair; $g_3$: fibroblast niche gene; $\chi^2$: test statistic for group-specific modulation of the dependence parameter.

| $g_1$ | $g_2$ | $g_3$ | $\chi^2$ | $g_1$ | $g_2$ | $g_3$ | $\chi^2$ |
| --- | --- | --- | --- | --- | --- | --- | --- |
| ACTN1 | ANXA3 | CFB | 39.70 | C1R | CXCL6 | GATM | 52.18 |
| ACTN1 | ANXA3 | GATM | 41.26 | ACTN1 | FLNA | AMY2A | 36.29 |
| CFB | FSTL3 | AMY2A | 106.89 | ACTN1 | FLNA | ANPEP | 31.13 |
| CFB | FSTL3 | ANPEP | 62.76 | ACTN1 | FLNA | CFTR | 31.39 |
| CFB | FSTL3 | BIRC3 | 35.62 | ACTN1 | FLNA | GATM | 54.54 |
| CFB | FSTL3 | CFTR | 38.13 | BIRC3 | CFTR | CTSK | 38.42 |
| CFB | FSTL3 | CTHRC1 | 34.67 | CXCL2 | FSTL3 | AMY2A | 47.94 |
| CFB | FSTL3 | CXCL6 | 32.76 | CXCL2 | FSTL3 | ANPEP | 44.46 |
| CFB | FSTL3 | GATM | 106.62 | CXCL2 | FSTL3 | CFB | 49.45 |
| CFB | FSTL3 | RNASE1 | 33.23 | CXCL2 | FSTL3 | CTSK | 31.01 |
| C1R | CFB | CTHRC1 | 37.21 | CXCL2 | FSTL3 | CXCL6 | 38.54 |
| CFB | PABPC1 | CFTR | 35.71 | CXCL2 | FSTL3 | GATM | 52.84 |
| CFB | PABPC1 | GATM | 33.90 | CXCL2 | FSTL3 | RNASE1 | 40.56 |
| ANPEP | FSTL3 | AMY2A | 33.71 | CFB | CXCL6 | AMY2A | 32.79 |
| ANPEP | FSTL3 | BIRC3 | 38.20 | CFB | CXCL6 | ANPEP | 56.96 |
| ANPEP | FSTL3 | CFB | 36.74 | CFB | CXCL6 | BIRC3 | 47.39 |
| ANPEP | FSTL3 | CFTR | 32.34 | CFB | CXCL6 | CFTR | 39.01 |
| ANPEP | FSTL3 | CTHRC1 | 31.74 | CFB | CXCL6 | GATM | 47.49 |
| ANPEP | FSTL3 | CTSK | 33.40 | CXCL2 | CXCL6 | CFB | 35.29 |
| ANPEP | FSTL3 | CXCL6 | 41.12 | CXCL2 | CXCL6 | GATM | 45.16 |
| ANPEP | FSTL3 | GATM | 32.32 | CXCL2 | CXCL6 | RNASE1 | 32.00 |
| C1R | CXCL6 | AMY2A | 31.60 | ACTN1 | RHOA | GATM | 32.88 |
| C1R | CXCL6 | ANPEP | 49.69 | CFB | SPIB | AMY2A | 33.99 |
| C1R | CXCL6 | CFB | 36.79 | CFB | SPIB | ANPEP | 36.68 |
| C1R | CXCL6 | CFTR | 33.98 | CFB | SPIB | GATM | 34.85 |
